# Etifoxine inhibits NLRP3 inflammasome activity in human and murine myeloid cells

**DOI:** 10.1101/2023.09.19.558428

**Authors:** Jordan M. Osmond, John B. Williams, Paul M. Matthews, David R. Owen, Craig S. Moore

## Abstract

**Background:** Multiple sclerosis (MS) is a chronic neuroinflammatory disease that is characterized by immune-mediated demyelination within the central nervous system. NLRP3 inflammasome activation has been previously reported as a possible pathophysiological contributor to microglial activation and oligodendroglial loss in MS, particularly in progressive forms of the disease.

**Methods:** Using both *in vivo* and *in vitro* approaches, this report investigated the use of a previously described ligand of the 18kDa translocator protein (TSPO), etifoxine, as an immunomodulator that inhibits inflammasome activation in primary human and murine macrophages and microglia. To further elucidate pathologic relevance in the MS context, investigations were also performed *ex vivo* using peripheral blood mononuclear cells and purified CD14^+^ monocytes derived from secondary progressive MS patients.

**Results:** Herein, it is demonstrated that etifoxine attenuated clinical symptoms in a mouse model of MS and significantly inhibited NLRP3 inflammasome activation in human and murine myeloid-derived cells *in vitro* by decreasing inflammasome-associated genes and inflammatory cytokine production. These anti-inflammatory effects of etifoxine were mediated independently of its previously described mechanisms related to engagement with TSPO and the GABA_A_ receptor. Furthermore, we observed a similar anti-inflammatory effect of etifoxine on MS patient-derived monocytes, which provides clinical relevance for the investigation of etifoxine as a potential therapeutic in progressive MS. Lastly, through the use of a gene array, we identified multiple signalling pathways in order to elucidate a novel mechanism whereby etifoxine may be inhibiting NLRP3 inflammasome activation.

**Conclusions:** Our results suggest that the anti-inflammatory effects of etifoxine were mediated independently of its previously described mechanisms related to engagement with TSPO and the GABA_A_ receptor. Furthermore, we observed an anti-inflammatory effect in murine- and human-derived myeloid cells, as well as in MS patient-derived monocytes, which provides clinically relevant evidence to support the exploration of etifoxine as a possible form of therapy for secondary progressive MS.

## BACKGROUND

Multiple sclerosis (MS) is a chronic demyelinating disease of the central nervous system (CNS) that is characterized by an infiltration of myelin-reactive CD4^+^ T cells (1, 2) and a disease pathology that consists of inflammation, axonal loss, and neurodegeneration within the brain and/or spinal cord. Clinically, MS is classified into three unique phenotypes or disease courses: 1) relapsing-remitting MS (RRMS), 2) secondary progressive MS (SPMS), and 3) primary progressive MS (PPMS) (2–5). Due to the heterogenous nature of MS, the cause(s) of disease prevalence, progression, and severity are presently unknown (6–8). In regard to treatment options, there are multiple disease-modifying therapies (DMTs) that are widely available to treat RRMS, however, only few options are available for patients with SPMS and PPMS (9, 10). This is largely due to the inability to fully understand the immune mechanisms related to the pathology of progressive forms of the disease; however, there is evidence to suggest that chronic microglial activation contributes to disease progression as a result of inflammasome-mediated activity (6, 9, 11–13).

Inflammasomes are molecular complexes that are present within immune cells comprising the innate immune system and are activated in response to cellular stress and pathogens. The NOD-like receptor family pyrin domain containing 3 (NLRP3) is the most widely recognized inflammasome and regulates caspase-1 activation, along with the maturation and production of IL-1β and IL-18 (1, 14–16). Inflammasome activation is induced by varying stimuli, such as reactive oxygen species and mitochondrial dysfunction, with the end result being a type of inflammasome-mediated cell death termed pyroptosis (17, 18). In MS, inflammasomes have been shown to be involved in disease progression and disease severity, specifically in progressive MS. Furthermore, inflammasomes and their associated signalling pathways have been identified as possible therapeutic targets for MS (18–21).

The 18kDa translocator protein (TSPO) is expressed in all cell types in the periphery, as well as the nervous system, and is located on the outer mitochondrial membrane where it is primarily involved in shuttling cholesterol, the rate-limiting step of steroid synthesis (22–24). TSPO has been identified as one of the most widely used targets for radioligands in positron emission tomography (PET) imaging, specifically in regard to the study of neuroinflammatory diseases (23, 25–27). In MS, chronic microglial activation has been linked to the enhanced neurodegeneration that occurs in progressive disease states; therefore, the use of TSPO-PET can aid in understanding the disease pathology of SPMS in relation to microglial activation, and may provide deeper insight into developing possible therapies for SPMS (19, 27–29).

Etifoxine was originally used clinically as a treatment for anxiety disorders, however, it has previously been identified as a TSPO ligand and also binds the ionotropic GABA_A_ receptors (30–32). In addition, etifoxine has also been reported to modulate inflammatory responses in both the central and peripheral nervous systems, albeit the mechanisms of action are presently unclear (33–35). Therefore, given the beneficial clinical effects of etifoxine in pre-clinical MS models (36, 37), we hypothesized that a novel mechanism of action for etifoxine may be related to its ability to modulate inflammasome activation, a critical immune-mediated pathological state driving oligodendroglial death in MS (38, 39).

In this study, we demonstrated that a previously described TSPO ligand, etifoxine, possesses anti-inflammatory properties that are mediated by the inhibition of NLRP3 inflammasome activation. Furthermore, we have demonstrated proof-of-principle that this novel mechanism of action may be therapeutically relevant in diseases, such as multiple sclerosis (MS) whereby heightened inflammasome activity is believed to be a pathological feature of disease progression.

## METHODS

### Mouse Primary Bone Marrow-Derived Macrophages

Primary bone marrow-derived macrophages (BMDMs) were isolated from C57BL/6 mice (Charles River). Mice aged 6 to 9 weeks were euthanized by CO_2_ followed by cervical dislocation. 70% ethanol was used to disinfect the body of the mice and the hind limbs were removed at the hip flexor. The muscle and adipose tissue were carefully cut from the bone; the cleaned bones were then placed in 50mL conical tubes containing phosphate buffered saline (PBS), which was then placed on ice. Separate conical tubes were used for each individual mouse preparation, and instruments were sterilized with 70% ethanol between each dissection. Femurs and tibias were then placed in petri dishes (one per animal) and further cleaned via the use of a scalpel. Once there was no tissue remaining on the bones, the femur and tibia were separated via a cut at the knee joint and the ankle joint. A scalpel was used to further cut the tops and bottoms of the individual bones to expose the bone marrow. The bone marrow was expelled from the bones using a 21½ gauge needle and 1mL of PBS. The initial 1mL of PBS was used to flush out the marrow from the remaining bones for each animal. Bone marrow was transferred to a 15mL conical tube and 3mL of ice-cold 0.8% ammonium chloride (StemCell) was added to lyse red blood cells. The 15mL conical was then placed on ice for 5 to 10 minutes. 10mL of PBS was then added to dilute the ammonium chloride and the conical was centrifuged at 450 x g for 10 minutes at 4°C. The supernatant was removed and the cell pellet was resuspended in 2mL of macrophage media consisting of DMEM (Thermofisher/Life Technologies), 10% heat inactivated fetal bovine serum (HI FBS) (USA Sourced, Corning, USA), 1x antibiotic/antimycotic (P/S) (Thermofisher/Life Technologies), 1x GlutaMAX (Thermofisher/Life Technologies), and 10ng/mL of Macrophage Colony-Stimulating Factor (M-CSF) (Cedarlane/Peprotech). 10µL of the cell suspension was then added to 90µL of trypan blue (Sigma Life Science) and a cell count was performed using a hemocytometer. Cells were plated at 5×10^5^ cells/mL in 10mL of macrophage media in a 10cm petri dish. After 3 days, an additional 5mL of macrophage media was added to the petri dish containing the cells. The cells were then plated in the experimental vessel after a total of 6-7 days. Plating macrophages involved removing the macrophage media that was currently on the cells and adding 5mL of ice-cold PBS. The petri dish was placed in the fridge for 15 minutes and a sterile cell lifter (Thermofisher/Life Technologies) was used to scrape the adhered cells from the bottom of the dish. The contents of the dish were transferred to a 15mL conical and centrifuged at 450 x g for 5 minutes. Cells were resuspended in 1mL of macrophage media (no M-CSF added) and recounted. The cell concentration was adjusted to 2.5×10^5^ cells/mL, and cells were plated in the experimental vessel. After 1-2 days in culture, the cells were ready to use for experimentation.

### Mouse Primary Microglia

The cortices of C57BL/6 mouse pups aged postnatal day 1 (P1) to postnatal day 4 (P4) were dissected to obtain a mixed glia culture. Prior to dissection, the following solutions were made: Dissection solution (DS) consisted of 1x sterile Hank’s Balanced Salt Solution (HEPES) (Thermofisher/Life Technologies) and HEPES (Sigma Life Science) with a final concentration of 10mM, which was then placed on ice for later use. Digestion solution consisted of 1mL 2.5% Trypsin (Thermofisher/Life Technologies), 2mL DS, and 0.05mg/ml DNAse 1 (Sigma Life Science); this volume of digestion solution is used for one culture (3 brains). Lastly, astrocyte media consisting of DMEM, 10% HI FBS, 1x P/S, and 1x GlutaMAX was prepared and pre-warmed in the 37°C water bath. Throughout the entire procedure, one culture is equivalent to three brains, and separate surgical instruments and solutions were used for each culture.

Mouse pups were euthanized by decapitation and the brain was removed and placed in a sterile petri dish that contained approximately 5mL of dissection solution (DS). Forceps were used to gently remove the meninges from the cortex and the cortices were isolated and transferred into another petri dish containing DS. Sterilized scissors were used to cut the cortices into smaller pieces and a transfer pipette was used to move the pieces of cortices to a 15mL conical tube. One 15mL conical tube contained the cortices that were isolated from three separate mouse brains. DS was carefully removed from the 15mL conical tube (~9mL), 3mL of digestion solution was added, and the contents of the tube were triturated using a transfer pipette. The tube was nutated in a 37°C incubator for a total of 15 minutes and gently triturated halfway through the incubation period. 500µL of HI-FBS was then added to stop the digestion (i.e. inactivate the trypsin) and the tube was placed back on the nutator for an additional 5 minutes. The sample was centrifuged at 200 x g for 5 minutes at room temperature. After centrifugation, the supernatant was removed and the sample was resuspended in 5mL of pre-warmed astrocyte media. The sample was initially triturated with a stuffed Pasteur pipette, followed by an 18-gauge blunt tipped needle, and lastly by a 21-gauge blunt tipped needle to obtain a single cell suspension. The cells were passed through a sterile 70 µm strainer into a 50mL conical and centrifuged at 300 x g for 5 minutes. The supernatant was removed, and the sample was resuspended in 1mL of pre-warmed media. Following trituration, the cell suspension was brought to 12mL with pre-warmed media and then transferred to a T75 flask and placed in a 37°C incubator. The following day, a full media change was performed with pre-warmed media and a two-thirds media change was performed on day 3 and again on day 6.

Microglia isolation occurred between days 10 and 14, depending on culture confluency. The standing incubator was pre-set to 37°C and the flask was shaken at 200rpm for 1 hour. The media from the flask was transferred to a 15mL conical tube and centrifuged at 400 x g for 10 minutes at room temperature. Microglia were resuspended in 1mL of astrocyte-conditioned media and 10µL of the cell suspension was added to 90µL of trypan blue for counting. Cells were then plated at 2.5 x 10^5^ cells/mL in the experimental vessel(s). The microglia were used 3-5 days after plating.

### 2.1 Experimental Autoimmune Encephalomyelitis

Experimental autoimmune encephalomyelitis (EAE) was induced in accordance with the protocols previously described (40, 41). 24 hours after first clinical signs (limp/flaccid tail), the mice were injected i.p. with vehicle (90% cyclodextrin, 10% ethanol), XBD-173 (10mg/kg twice daily), or etifoxine (50mg/kg once daily, saline once daily) (Sigma Aldrich) for 7 consecutive days. After 7 days of treatment, the mice were euthanized using sodium pentobarbital, transcardially perfused with sterile saline, and followed by cervical dislocation. Brains and spinal cords were removed for subsequent macrophage/microglia isolation (42). Briefly, tissues were homogenized in RPMI using a dounce homogenizer to obtain a single cell suspension. Using a percoll gradient and various centrifugations and washes, the cells were collected at the 70%-30% gradient interface and MACS was performed on these cells using CD11b+ beads to obtain a pure population of microglia according to manufacturer’s protocols (Miltenyi). Cells were then stored in Trizol at −80°C.

### Human Primary Monocyte-Derived Macrophages

Human monocyte-derived macrophages (MDMs) were cultured from venous whole blood. All studies that involved human samples followed Canadian Institute of Health Research (CIHR) guidelines and institutional review board approval at Memorial University of Newfoundland (Health Research Ethics Authority). Peripheral blood was collected from healthy donors in EDTA-coated tubes following informed consent. Blood was placed on the nutator at room temperature while preparing for cell isolation. The following solutions were made: Macrophage media consisted of RPMI 1640, 10% HI-FBS, 1x P/S, 1x GlutaMAX, and 25ng/mL of M-CSF. MACS buffer was used for cell separation and consisted of 500mL sterile 1x PBS, 2mL of 0.5M EDTA, and 2.5mL of HI-FBS. Blood was pooled from two 10mL BD Vacutainer® tubes into a 50mL conical tube. Tubes were then rinsed with sterile PBS and the remaining contents were emptied into the 50mL conical tube. The conical tube was filled with 1x PBS to a total volume of 35mL. SepMate™ tubes (StemCell Technologies) were used to isolate peripheral blood mononuclear cells (PBMCs). 15mL of Ficoll-Hypaque (ThermoFisher/Life Technologies) was added directly through the hole in the SepMate™ tube, next the 35mL blood/PBS mixture was slowly added to the tube via the use of a 25mL serological pipette. The SepMate™ tube was centrifuged at 1200 x g for 10 minutes (brake on) to separate the contents of the tube. The entire top layer (~35mL) was poured into a 50mL conical, which was then filled to 50mL with 1x PBS. The cells were centrifuged at 300 x g for 15 minutes. The supernatant was poured off and the pellet was resuspended in 20mL of MACS buffer; an aliquot was taken and diluted at 1:2 in Trypan Blue to determine the number of cells. The sample was then brought to 50mL with MACS buffer and centrifuged at 300 x g for 10 minutes. The supernatant was poured off and the cells were resuspended in 80µL of MACS buffer/1×10^7^ cells. 20µL of anti-CD14^+^ (Miltenyi Biotec)/1×10^7^ cells was also added to the sample, which was then mixed and incubated at 4°C for 15 minutes. During the incubation period, the MACS column was rinsed with 3mL of MACS buffer. After 15 minutes, the sample was washed with 20mL of MACS buffer and centrifuged at 300 x g for 10 minutes. The supernatant was poured off and the cells were resuspended up to 1×10^7^ cells/mL of MACS buffer. The cell suspension was poured into the column and flow through was collected in a 15mL conical tube. The column was washed with 3mL of MACS buffer 3 consecutive times; the flow through was discarded. The column was removed from the magnet and placed on a new 15mL conical tube. 5mL of MACS buffer was added to the column and quickly plunged to expel the CD14+ antibody-bound cells. A cell count was conducted, the final volume was brought to 15mL with MACS buffer, and the sample was centrifuged at 300 x g for 10 minutes. Cells were then resuspended in macrophage media and plated in desired experimental vessels at 5×10^5^ cells/mL. After 2-3 days a half media change (RPMI 1640, 10% HI-FBS, 1x P/S, 1x GlutaMAX, and 25ng/mL of M-CSF) was performed, and the cells were used for experimentation 3-4 days later.

### Human Primary Fetal Microglia

Human microglia were isolated from fetal CNS tissue obtained from consenting donors (Health Sciences Centre – General Hospital, St. John’s, NL). Prior to dissecting the culture media was made and pre-warmed in a 37°C water bath. Culture media consisted of DMEM, 5% HI-FBS, 1x P/S, and 1x GlutaMAX. The CNS tissue dissection was performed using a stereomicroscope and sterilized surgical instruments. Sections of the brain and/or spinal cord were isolated and transferred to a 15mm petri dish containing sterile saline. The meninges were removed via the use of sterile forceps, and the tissue was transferred to a 15mL along with 2mL of sterile 1x PBS, 1mL of 2.5% Trypsin, and 200µL of DNase. The sample was triturated with a 1000µL pipette and was then incubated at 37°C for 15 minutes on a nutator. Following the incubation period, 1mL of HI-FBS was added to the sample, which was then triturated and passed through a 70µm filter into a 50mL conical. The filter was rinsed with 1x PBS and the sample was centrifuged at 300 x g for 10 minutes at 4°C. The supernatant was removed, the sample was resuspended in 5mL of warmed culture media and then centrifuged at 300 x g for 10 minutes. Following centrifugation, the supernatant was removed, and the sample was resuspended in 4mL of culture media, which was then transferred to a T12.5 flask. Fetal microglia were isolated and cultured according to (43).

### HITMS Patient-Derived Primary Monocyte-Derived Macrophages

All participants had provided written consent to be enrolled in the HITMS project at Memorial University of Newfoundland. The patient demographics have been described in Table 4. All studies that involved human samples followed Canadian Institute of Health Research (CIHR) guidelines and institutional review board approval at Memorial University of Newfoundland (Health Research Ethics Authority).

PBMCs were isolated from whole blood in accordance with the protocol described above, after performing the cell count monocytes were isolated from PBMCs or whole-PBMCs were cryopreserved. Monocyte (CD14+ cell) isolation was performed according to the MACS separation protocol above. To differentiate the monocytes into macrophages, media comprised of RPMI, 10% HI-FBS, 1x P/S, 1x GlutaMAX, and 25ng/mL of M-CSF was used; To culture monocytes, similar media was used, however, no M-CSF was added and the cells were cultured upright in a 15mL conical tube with a vented cap.

Cryopreservation was conducted by resuspending the cells in ice-cold PBMC storage media, which consisted of 80% HI-FBS and 20% Culture Media (RPMI, P/S, 1x GlutaMAX). An equal volume of ice-cold freezing media, which consisted of 70% HI-FBS, 10% Culture Media, and 20% DMSO, was slowly added to the sample and gently mixed. 1.0 mL aliquots of the cell suspension were then distributed into cryovials that were placed in a room temperature cryofreezing container, which was then stored at −80°C overnight. The cryovials were then transferred to liquid nitrogen for long-term storage. The patient information and corresponding cell count were updated in the HITMS database.

### Inflammasome Activation and Etifoxine Treatment

Inflammasome activation was conducted in the various cell types described above. Initially, a media change was performed on the cells that were being treated, thus the following protocol was conducted in a serum-free environment (DMEM or RPMI, 1x P/S, 1x GlutaMAX). Once in serum-free media, cells were pre-treated with etifoxine (500nM to 5000nM) (10mM stock prepared in DMSO and stored at −20°C)) for a total of 1 hour. The dose of etifoxine was cell type dependent, mouse-derived cells were pre-treated with 0.5uM – 5uM etifoxine and human-derived cells were pre-treated with 5uM – 50uM etifoxine. To determine whether the effect of etifoxine was mediated via TSPO, cells were simultaneously treated with etifoxine and established TSPO ligands (XBD173 and PK11195) at doses well-above the established K_d_ (44, 45) After 1 hour of pre-treatment, lipopolysaccharide (LPS) (100ng/mL) was pipetted directly into the wells. The cells were primed with LPS for 3 hours and then activated with nigericin (10µM) for 1 hour. Following activation, the supernatants were collected in 1.5mL Eppendorf tubes, and stored at −80°C. The adherent cells that remained were either lysed with standard radioimmunoprecipitation assay (RIPA) buffer with 1mM Na_3_VO_4_ and BD Baculogold protease inhibitors (BD Biosciences) or 500uL of QIAzol® reagent (Qiagen). Samples were then transferred to 1.5mL Eppendorf tubes and stored at −80°C.

Inflammasome activation and etifoxine treatment were also conducted in HITMS patient-derived whole-PBMCs. All patients were diagnosed with secondary progressive MS and were not presently using any disease modifying therapies (DMTs). Only patient samples that exhibited a positive response following inflammasome activation (IL-1β response measured via ELISA) were included. Patient demographics are outlined in Table 4.

Prior to thawing PBMCs, culture media (RPMI, P/S, 1x GlutaMAX) was warmed to 37°C. PBMCs were thawed by immersing cryovials into a 37°C water bath, however vials were not fully immersed to ensure that water did not seep through the cap and contaminate the sample. Once the PBMCs were mostly thawed (small ice crystals present in cryovial), the 1.0mL cell suspension was transferred to a 15mL conical. 9.0mL of pre-warmed media was then added to the 15mL conical containing the PBMC cell suspension. Samples were centrifuged at 300 x g for 10 minutes to obtain a cell pellet. The supernatant was carefully poured off and the cells were resuspended in 1mL of pre-warmed media, an aliquot was taken and diluted at 1:2 in Trypan Blue to determine the number of viable cells. Cells were then resuspended at 5×10^5^ cells/mL with pre-warmed media in 5mL polypropylene round-bottom tubes. After 1-2 days the samples were centrifuged at 300 x g for 10 minutes and resuspended in serum-free media. Cell suspensions were then pre-treated with etifoxine (50uM) and inflammasome activated according to the protocol outlined above. Following the inflammasome activation protocol the cell suspensions were transferred to 1.5mL Eppendorf tubes and micro-centrifuged at 300 x g for 10 minutes. The supernatants were aliquoted into new 1.5mL Eppendorf tubes and stored at −80 °C and the cell pellets were flash frozen in liquid nitrogen and stored at −80 °C.

### Picrotoxin Treatment

BMDMs and THP-1 macrophages were plated in 24-well plates and pre-treated with 50uM picrotoxin alone or with 5uM – 10uM etifoxine. Following the 30-minute pre-treatment, the cells underwent inflammasome activation in accordance with the protocol described above. Supernatants were collected, transferred to 1.5mL Eppendorf tubes, and stored at −80°C.

### RNA Isolation, Reverse Transcription, and Quantitative Polymerase Chain Reaction (RT-qPCR)

The RNeasy Micro Kit (Qiagen) was used to extract RNA from cells that were lysed with QIAzol® reagent (Qiagen). The extraction was conducted according to the manufacturer’s protocol. The concentration of RNA was determined by using a Nanodrop 1000 Spectrophotometer (Fisher Scientific). Reverse transcription was performed by using the High Capacity cDNA Reverse Transcription Kit (Applied Biosystems) according to the manufacturer’s protocol. Polymerase chain reaction (PCR) was conducted with the Taqman® Fast Universal PCR Master Mix (Applied Biosystems). The TaqMan® probes and primers that were used are as follows: *mIL-1β*, *hIL-1β, mNLRP3, hNLRP3, mCaspase-1*, *18S Endogenous Control,* and *hGAPDH Control* (Applied Biosystems). RT-qPCR was performed using an Applied Biosystems® ViiA 7 Real-Time PCR System and analysis was conducted on QuantStudio^TM^ Software by Applied Biosystems. Minus reverse transcriptase and no-template cDNA controls were included. Fold changes were calculated using the 2^-ΔΔCT^ method(46).

### RT^2^ Profiler PCR Array

RNA extraction and quantification were conducted according to the procedure(s) stated above. Reverse transcription was performed by using the RT^2^ First Strand Kit (Qiagen) according to the manufacturers protocol. PCR was conducted with the Mouse Inflammasomes (96-well format) RT^2^ Profiler PCR Array (Cat. # PAMM-097Z, Qiagen) according to the manufacturers protocol and was read on an Applied Biosystems® ViiA 7 Real-Time PCR System and analysis was conducted on Quantstudio^TM^ Software by Applied Biosystems. The layout of the PCR array 96-well plate consisted of probes/primers for 84 inflammasome-associated genes, 5 housekeeping controls, 1 mouse genomic DNA contamination control, 3 reverse transcription controls, and 3 positive PCR controls. C_T_ values for each 96-well plate were exported to a Microsoft Excel^®^ spreadsheet, which was then uploaded to the Qiagen GeneGlobe Data Analysis Center online-based software for further analyses, with instructions provided in the manufacturers protocol.

Genes were considered to be up or downregulated when the fold change was greater than 2, which is represented by the blue lines on the scatterplots. The fold change cut off could be changed, however the default option was set at 2. Genes were considered to be unchanged (black) when they were below the fold change cut off value.

### Enzyme-Linked Immunosorbent Assay

Human IL-1β, tumor necrosis factor (TNF), IL-6, and mouse TNF OptEIA enzyme-linked immunosorbent assay (ELISA) kits were used from BD Biosciences. Human IL-18 and mouse IL-1β ELISA kits were used from R&D Systems (biotechne). All ELISAs were performed according to the manufacturer’s instructions. Absorbance readings were obtained by using a Cytation™ 5 Cell Imaging Multi-Mode Reader (BioTek) and concentrations were determined based on a standard curve with a linear line of best fit.

### Western Blotting

Human macrophages were inflammasome-activated according to the protocol above. Cells were lysed with radioimmunoprecipitation assay (RIPA) buffer and stored at −80°C. Prior to SDS-PAGE, protein lysates were thawed and diluted at a 1:1 ratio in 2X sample buffer and heated to 95°C for 5 minutes. Samples were separated on WedgeWell™ 4–15% Tris-Glycine Gels (Invitrogen, Burlington, Canada) and transferred to 0.45 pore Immobilon-P PVDF membranes (Millipore) for 1 hr at 100 V. Membranes were blocked overnight at 4°C in 5% skim milk in 1x tris-buffered saline and tween 20 (TBST). Membranes were then probed with antibodies specific to NLRP3 (1:1000) or β-actin (1:1000) followed by HRP-linked anti-IgG (1:2000). Membranes were then soaked in ECL™ Western Blotting Detection Agents and developed. Protein loading was normalized relative to β-actin.

### Propidium Iodide Uptake and Cytotoxicity Assay

Propidium iodide (ThermoFisher/Life Technologies) is a cell impermeable DNA stain that was used to mark cells that were undergoing inflammasome-mediated cell death (pyroptosis). The assays were conducted in 96-well plates, in which cultured BMDMs and MDMs were supplemented with 100µL/well serum-free and phenol-red free DMEM or RPMI. One column of the 96-well plate did not contain cells, so it was used to account for the background fluorescence. The background fluorescence reading for each row was subtracted from the corresponding wells that contained Triton X-100 (Sigma Aldrich) and the sample being measured; this was done to normalize the fluorescence index throughout the experimental reading.

For the propidium iodide uptake assay, inflammasome activation and etifoxine treatments were conducted according to the procedure above, up until the addition of nigericin. Prior to activating cells with nigericin, the serum-free media was replaced with serum-free and phenol-red free media that contained 1ug/mL of propidium iodide. At this point there were three different tubes containing media with propidium iodide: a no treatment condition, a nigericin (10uM) condition, and a Triton X-100 (10µL/mL) condition. The 96-well plate was then transferred to the Cytation™ 5 Cell Imaging Multi-Mode Reader (BioTek), which was set at 37°C +/- 2°C with 5% CO_2_. The fluorescence intensity was read at 533/617 (excitation/emission) every 2 minutes for 1.5 hours. The relative propidium iodide uptake was determined with the following equation: 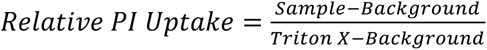.

For the cytotoxicity assay, the cells were only treated with various doses of etifoxine, which were all diluted in serum-free and phenol-red free media that contained 1ug/mL of propidium iodide. In BMDMs, the conditions included: no treatment, etifoxine (1uM, 3uM, 5uM, 10uM, 15uM, 30uM, 100uM, and 300uM), and Triton X-100. In MDMs the conditions were no treatment, etifoxine (1uM, 3uM, 10uM, 30uM, 100uM, and 300uM), and Triton X-100. The 96-well plate was then transferred to the Cytation™ 5 Cell Imaging Multi-Mode Reader, which was set at 37°C +/- 2°C with 5% CO_2_. The fluorescence intensity was read at 533/617 (excitation/emission) every 5 minutes for 12 hours. Instead of measuring propidium iodide uptake, the results were displayed as mean fluorescent units to better observe if there was a cytotoxic effect in any of the etifoxine conditions.

### Statistical Analysis

Statistical analyses were performed using GraphPad Prism Version 8.4.0. Data is presented in the form of the mean +/- standard error of the mean (SEM). Ordinary and repeated measures one-way analyses of variance (ANOVA) were conducted and the recommended post hoc tests that were used are mentioned in the corresponding figure descriptions. *p*<0.05 was considered significant.

## RESULTS

### Treatment with, etifoxine, but not XBD173, decreases clinical severity of EAE

After presenting with the first clinical signs of EAE (limp/flaccid tail), mice were administered either XBD173, etifoxine, or vehicle (24hrs after first clinical signs) to determine if these pre-established TSPO ligands were capable of decreasing clinical severity. Mice treated with etifoxine presented with less severe clinical symptoms compared to the vehicle and XBD173 treatment groups. The etifoxine-treated group (Supplemental Figure 1D) peaked at a mean score of 1.75 on day 6, which dropped to a mean score of 1.25 on day 8. In comparison, the vehicle (Supplemental Figure 1B) and XBD173 (Supplemental Figure 1C) groups peaked at a mean score of 2.75 and 2.66 on days 7 and 6, respectively.

### Etifoxine pre-treatment decreases IL-1β secretion in mouse-derived primary macrophages and microglia under inflammasome-activating conditions

To further elaborate on the mechanism by which etifoxine may be inhibiting the inflammatory response in EAE mice, macrophages generated from monocytes (Figure 1A) and primary microglia (Figure 1B) were cultured and treated with LPS and nigericin to activate the NLRP3 inflammasome, which has recently been suggested to be involved in driving the pathogenesis of MS. In the absence of inflammasome activation, etifoxine (5uM) alone did not have an effect on IL-1β secretion; 5uM dose was pre-determined based on literature involving cell culture work done in microglia and astrocytes as well as preliminary results. A significant decrease in IL-1β expression (*p*<0.001) was observed in the etifoxine (5uM) pre-treatment group in macrophages and microglia when compared to inflammasome only controls in both cell types.

**Figure 1:**
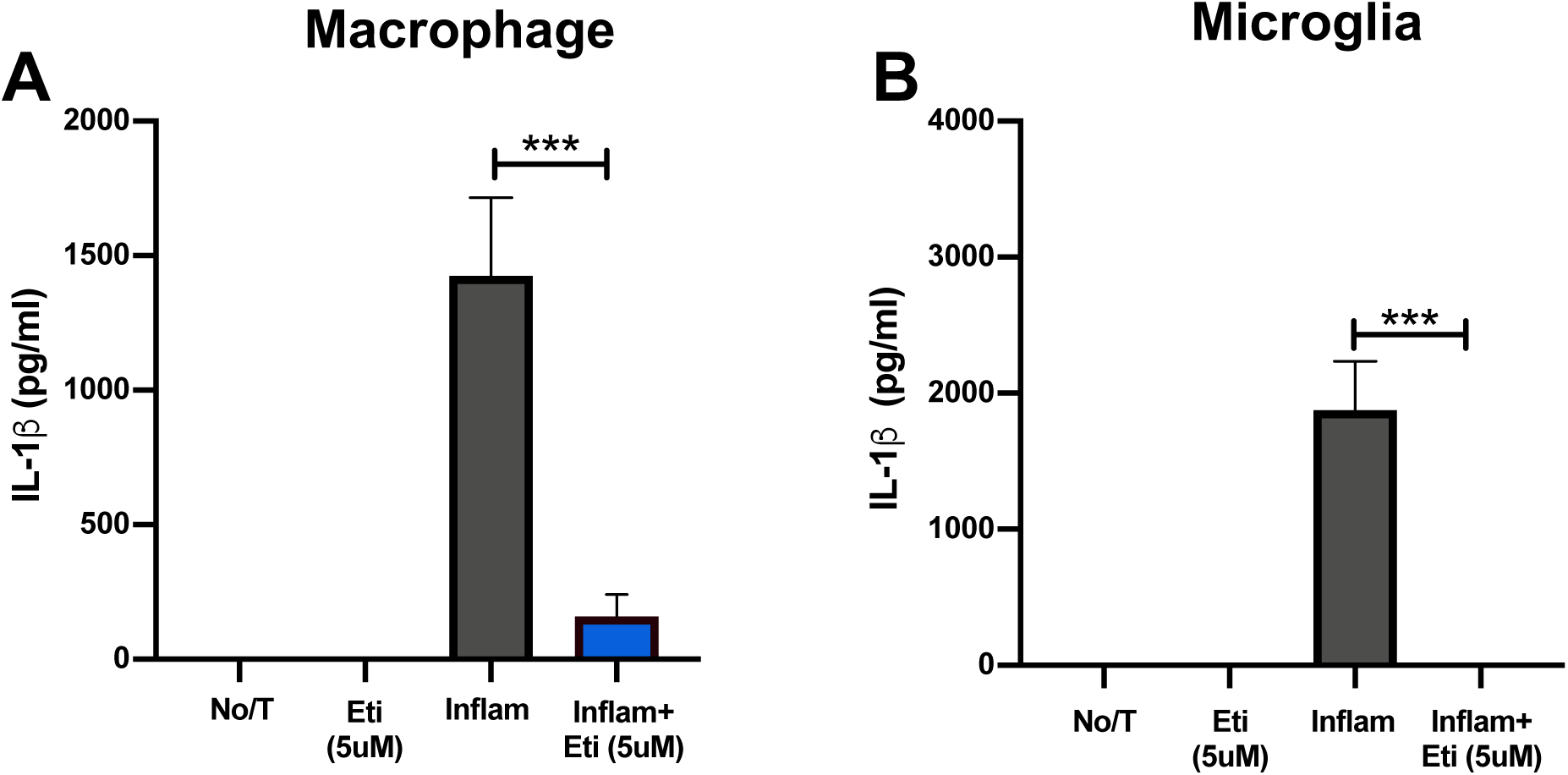
IL-1β secretion is significantly decreased in primary mouse macrophages and microglia when pre-treated with etifoxine (5uM) under inflammasome-activating conditions. IL-1β ELISAs were used to measure cytokine expression in supernatants of primary mouse macrophages (n=7) and microglia (n=8) that were either untreated, treated with etifoxine (5uM) (eti(5uM)), inflammasome activated (inflam), or pre-treated with etifoxine (5uM) under inflammasome-activating conditions (inflam + eti(5uM)). (A) IL-1β secretion in primary mouse macrophages was decreased when treated with etifoxine under inflammasome-activating conditions (160.2pg/mL±13.46) compared to the inflammasome-only control (1426pg/mL±100.30). (B) IL-1β secretion in primary mouse microglia was decreased when treated with etifoxine under inflammasome-activating conditions (0pg/mL) compared to the inflammasome-only control (1874pg/mL±146.32). Results are displayed as mean ± SEM. One-way analysis of variance with Tukey’s post hoc test was used to determine group differences. \*\*\**p*<0.001.

### Etifoxine shows dose-dependent inhibition of IL-1β secretion from human-derived primary macrophages and fetal microglia under inflammasome-activating conditions

To determine if the inhibitory effects of etifoxine were also translatable to humans, primary human macrophages and fetal microglia were isolated and activated in culture. A range of doses of etifoxine (0.1-50uM) were used in macrophages to determine an optimal dose. In macrophages (Figure 2A) a dose-dependent effect was observed; a decrease in IL-1β secretion was observed only with pre-treatment at concentrations > 25uM compared to the inflammasome-only control. IL-1β secretion decreased at the highest etifoxine concentration (50uM) when compared to the inflammasome-only control, but did not reach statistical significance.

**Figure 2:**
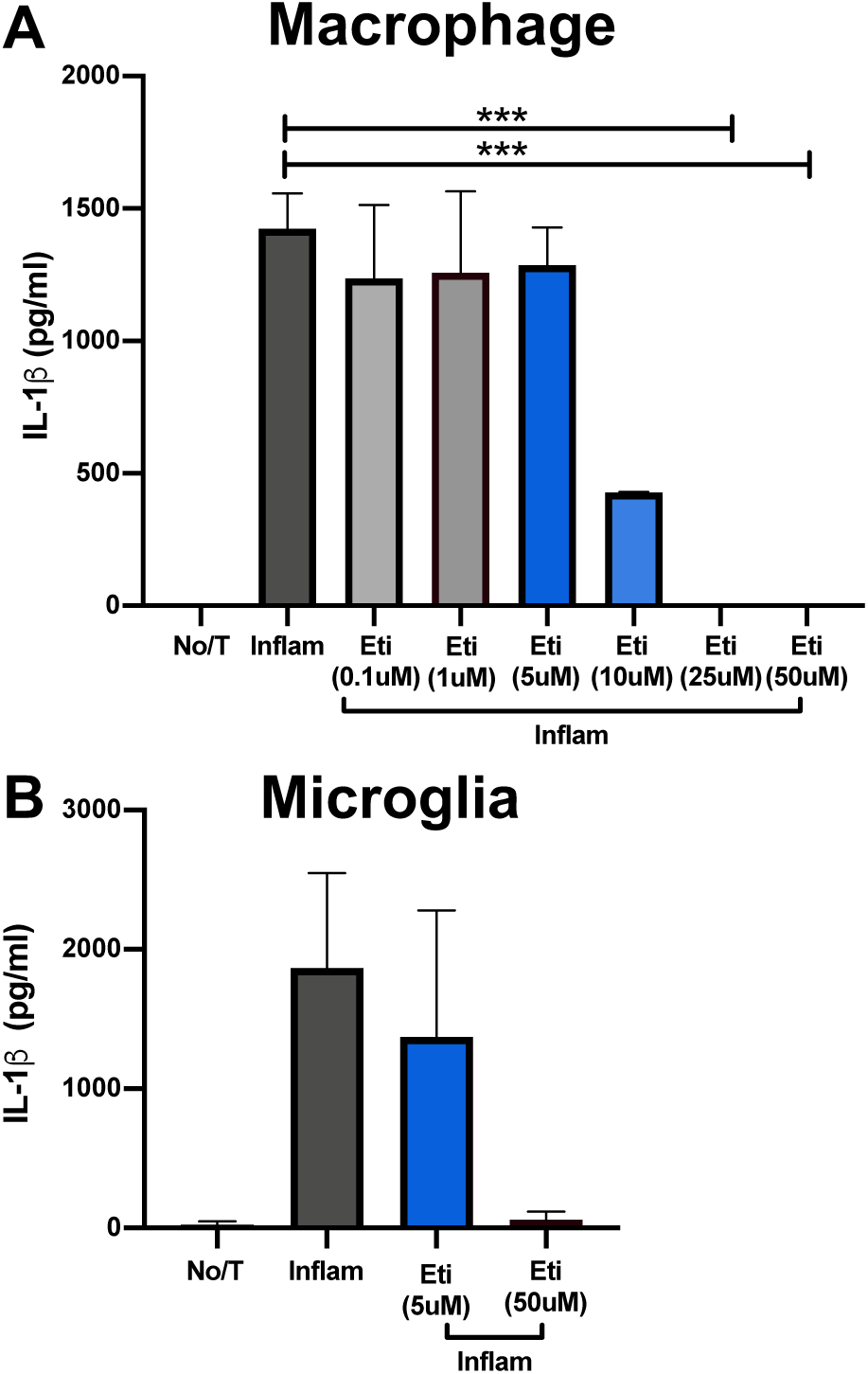
IL-1β secretion is significantly decreased in primary mouse macrophages and microglia when pre-treated with etifoxine (5uM) under inflammasome-activating conditions. IL-1β ELISAs were used to measure cytokine expression in supernatants of primary mouse macrophages (n=7) and microglia (n=8) that were either untreated, treated with etifoxine (5uM) (eti(5uM)), inflammasome activated (inflam), or pre-treated with etifoxine (5uM) under inflammasome-activating conditions (inflam + eti(5uM)). (A) IL-1β secretion in primary mouse macrophages was decreased when treated with etifoxine under inflammasome-activating conditions (160.2pg/mL±13.46) compared to the inflammasome-only control (1426pg/mL±100.30). (B) IL-1β secretion in primary mouse microglia was decreased when treated with etifoxine under inflammasome-activating conditions (0pg/mL) compared to the inflammasome-only control (1874pg/mL±146.32). Results are displayed as mean ± SEM. One-way analysis of variance with Tukey’s post hoc test was used to determine group differences. \*\*\**p*<0.001.

### Etifoxine inhibits IL-1β secretion when added either before or during LPS priming of inflammasome activation

Inflammasome activation *in vitro* occurs in 2 steps (LPS priming and nigericin execution), we sought to determine a more defined mechanism of action and when etifoxine may be exerting its anti-inflammatory effect during this activation protocol. To determine when etifoxine may be having its greatest anti-inflammatory effect, cells were either treated with etifoxine 30 minutes prior to LPS priming, during LPS priming, or during inflammasome execution with nigericin. The greatest decrease in IL-1β secretion occurred in the etifoxine (5uM) pre-treatment group and the simultaneous etifoxine (5uM) and LPS treatment group when compared to inflammasome-only control (Figure 3).

**Figure 3:**
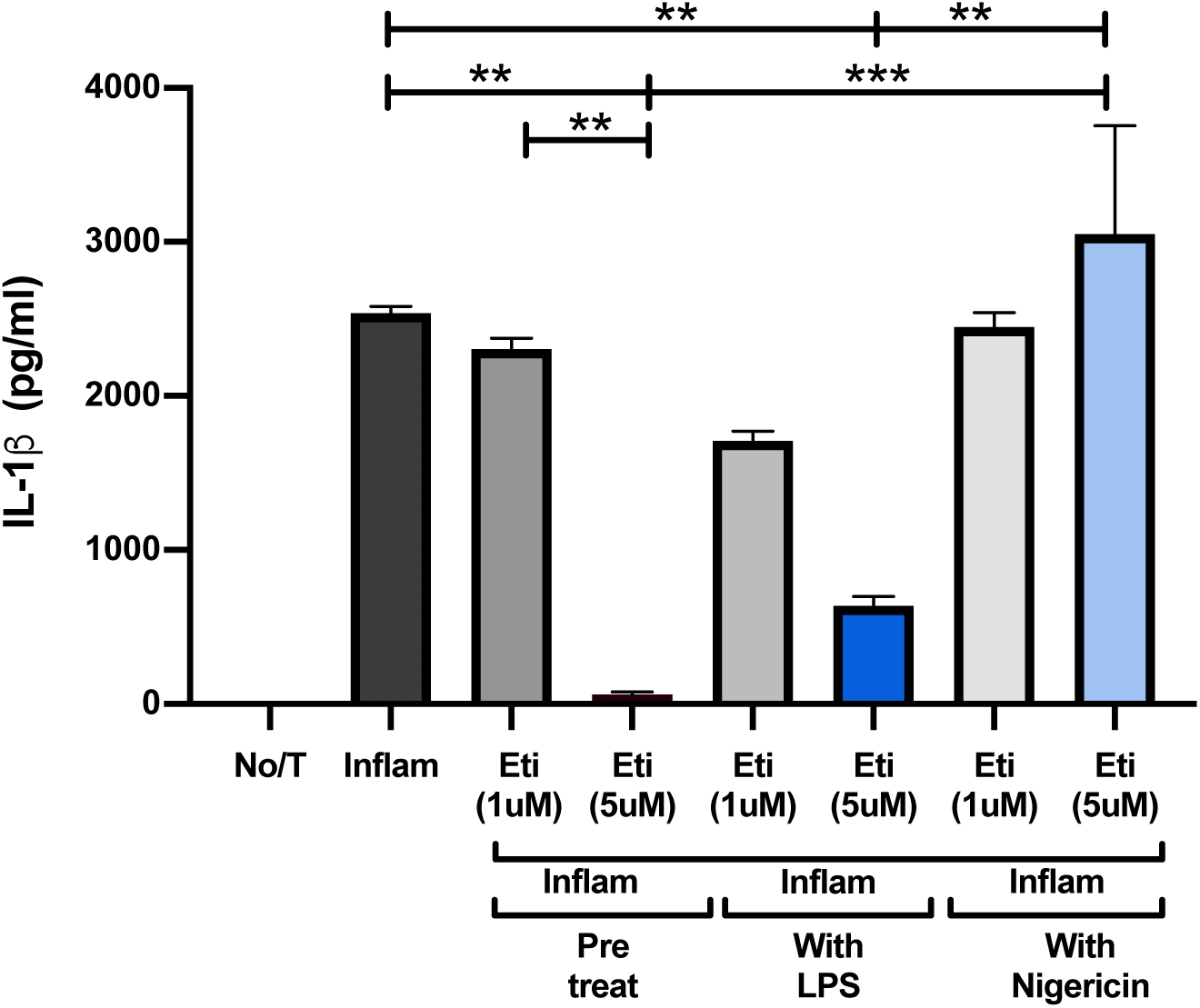
IL-1β secretion is decreased in primary mouse BMDMs that are either pre-treated with etifoxine (5uM) or when simultaneously treated with LPS. IL-1β secretion measured by ELISA in primary mouse macrophages (n=2) that were untreated, inflammasome activated (inflam) (2538pg/mL±117.62), and pre-treated with etifoxine (1uM, 5uM) (eti) prior to inflammasome activation (2305pg/mL±143.65, 57.21pg/mL±7.01), with LPS (1708pg/mL±59.055, 637pg/mL±19.92), or with nigericin (2450pg/mL±91.96, 3052pg/mL±148). Results are displayed as mean ± SEM. One-way analysis of variance with Tukey’s post hoc test was used to determine group differences. \*\**p*<0.01, \*\*\**p*<0.001.

### Under inflammasome-activating conditions, etifoxine decreases TNF release in human macrophages

LPS treatment alone will result in the release of TNF. To determine whether etifoxine was exerting its anti-inflammasome effects at either the priming and/or executing step of the inflammasome activation paradigm, TNF levels were measured in the supernatants. In the priming step (LPS activation), if levels of TNF were lower with etifoxine pre-treatment, it would suggest that etifoxine is inhibiting the LPS-TLR4 mediated signaling cascade, hence ultimately leading to less inflammasome activation once treated with the executing nigericin step. Under inflammasome-activating conditions, TNF secretion was significantly decreased (*p*<0.05, *p*<0.01) in human macrophages in a dose-dependent manner when pre-treated with etifoxine; (Figure 4). Alternatively, potassium influx can stimulate inflammasome activation. In murine macrophages (Figure 4) simultaneous treatment with etifoxine and nigericin did not affect inflammasome activation.

**Figure 4:**
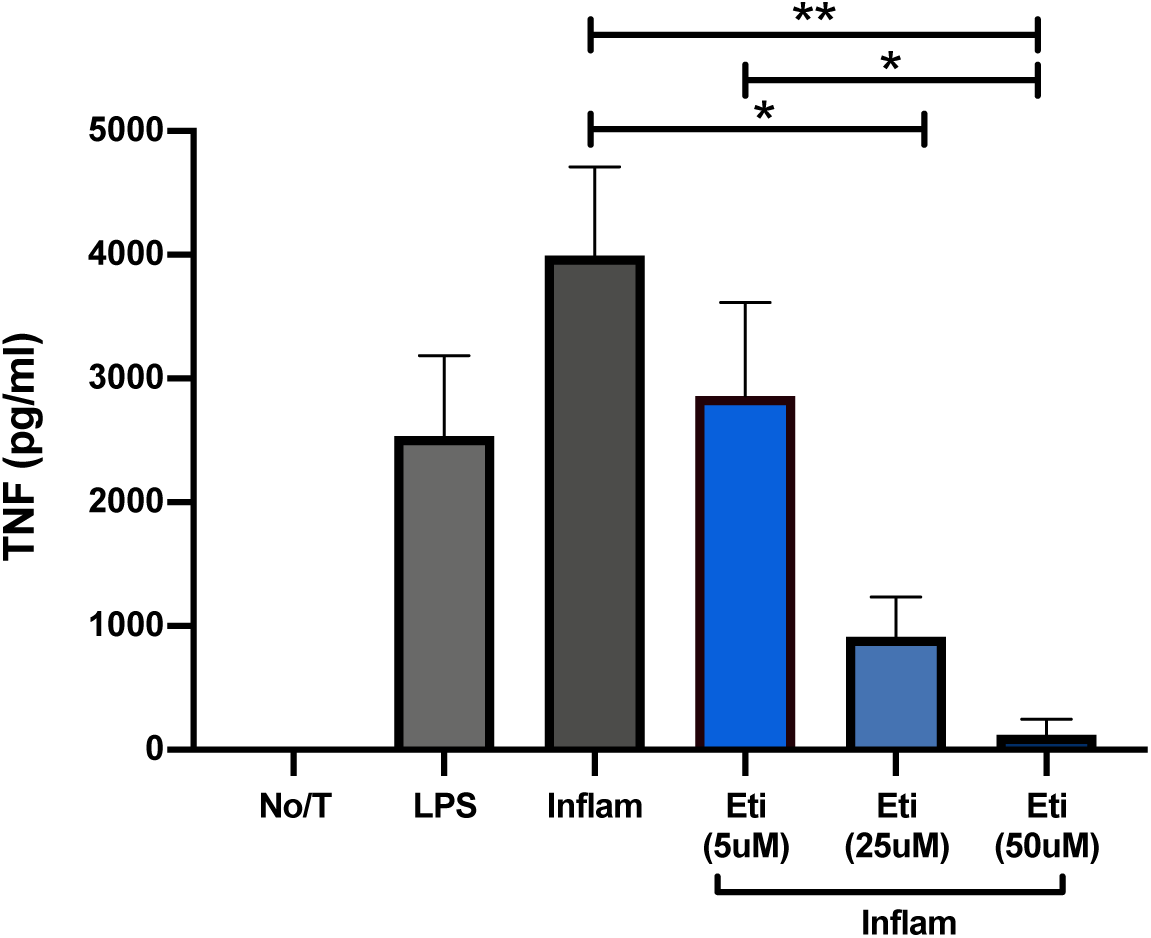
TNF secretion is decreased when primary human macrophages are pre-treated with etifoxine under inflammasome-activating conditions. A TNF ELISA was conducted on primary human macrophages (n=3) that were untreated, treated with LPS, inflammasome activated (inflam), and pre-treated with etifoxine (5,25,50uM) (eti) under inflammasome-activating conditions. IL-1β secretion significantly decreased in a dose-dependent manner when pre-treated with etifoxine (25uM,50uM) under inflammasome-activating conditions (913.95+/-42.71, 123+/-9.16) compared to the inflammasome-only control. Results are displayed as mean with standard error. One-way analysis of variance with Tukey’s post hoc test was used to determine group differences. \**p*<0.05, \*\**p*<0.01

### Etifoxine is not cytotoxic to naïve or inflammasome-activated mouse-derived and human-derived primary macrophages

A wide dose (1-300uM) cytotoxicity assay was conducted in naïve murine- and human-derived primary macrophages to determine if high concentrations of etifoxine were implementing a cytotoxic effect. In both human and murine macrophages, etifoxine did not increase the PI fluorescent signal when measured over a 5-hour time period, regardless of concentration (Supplemental Figure 2). Treatment with Triton X served as the positive control and quickly resulted in a positive PI signal that peaked at approximately 2700 MFU followed by a plateau in mice; approximately 2600 MFU in human cells. The PI signal in the etifoxine conditions did not exceed 900 MFU in either human or murine cells. This assay was also conducted to ensure that etifoxine was not initiating any form of cell death prior to the cells undergoing pyroptosis. Etifoxine alone did not affect cell viability (Supplemental Figure 3), so we wanted to determine if etifoxine treatment under inflammasome-activating conditions could rescue macrophages from undergoing pyroptosis. In murine macrophages, pre-treatment with etifoxine did not have an effect on PI uptake under inflammasome-activating conditions, this observation was present in both the untreated (mean difference of 0.03 or 3%) and inflammasome-only controls (mean difference of 0.03 or 3%). In human macrophages, PI uptake was decreased when pre-treated with etifoxine (50uM) under inflammasome-activating conditions in comparison to the inflammasome-only control (mean difference of 0.31 or 31%), and unchanged when compared to the untreated control (mean difference of 0.08 or 8%). This provided validation that etifoxine was not initiating a premature cell death under inflammasome-activating conditions.

### Etifoxine inhibition of inflammasome activation is not mediated by GABA_A_ receptors or TSPO

Etifoxine has previously been described to bind the GABA_A_ receptor, which has been demonstrated to be expressed on myeloid cells (47, 48). In regard to inflammation, prior findings have indicated that GABA_A_ receptors are capable of modulating inflammation, however the anti-inflammatory effect is highly dependent on the pharmaceutical agonist being used(49). In order to elucidate whether etifoxine might be working via a GABA_A_-mediated mechanism to inhibit inflammasome activation, picrotoxin, the prototypic GABA_A_ antagonist was used to block GABA_A_ receptors. In primary mouse macrophages (Supplemental Figure 4), pre-treatment with picrotoxin alone did not affect IL-1β secretion under inflammasome-activating conditions. In further investigation using THP-1 macrophages, picrotoxin did not influence IL-1β secretion (data not shown) and compared to the etifoxine-treated (5uM-10uM) inflammasome condition (levels were not statistically different. These results suggest that the ability of etifoxine to inhibit the inflammasome is not via the blockade of GABA_A_ receptors. In addition, the inability of established TSPO ligands (e.g. PK11195 and XBD173) to inhibit inflammasome activation at concentrations high above the their established K_d_ (44, 45) (either separately or in combination with etifoxine) suggest the mechanisms by which etifoxine is inhibiting the inflammasome is independent of its ability to bind TSPO (Supplemental Figure 5).

### Etifoxine pre-treatment downregulates inflammasome-associated genes in primary mouse-derived macrophages and microglia

RT-qPCR showed etifoxine pre-treatment in murine myeloid cells reduced expression of inflammasome-related genes, including *il-1β* and *nlrp3*. In macrophages, there was a 180-fold decrease in *il-1β* expression (Figure 5A) and a 112-fold decrease in *nlrp3* expression (Figure 5C) when compared to inflammasome-only controls. In microglia there was a 1125-fold decrease in *il-1β* expression (Figure 5B) and a 48-fold decrease in *nlrp3* expression (Figure 5D) when compared to inflammasome-only controls.

**Figure 5:**
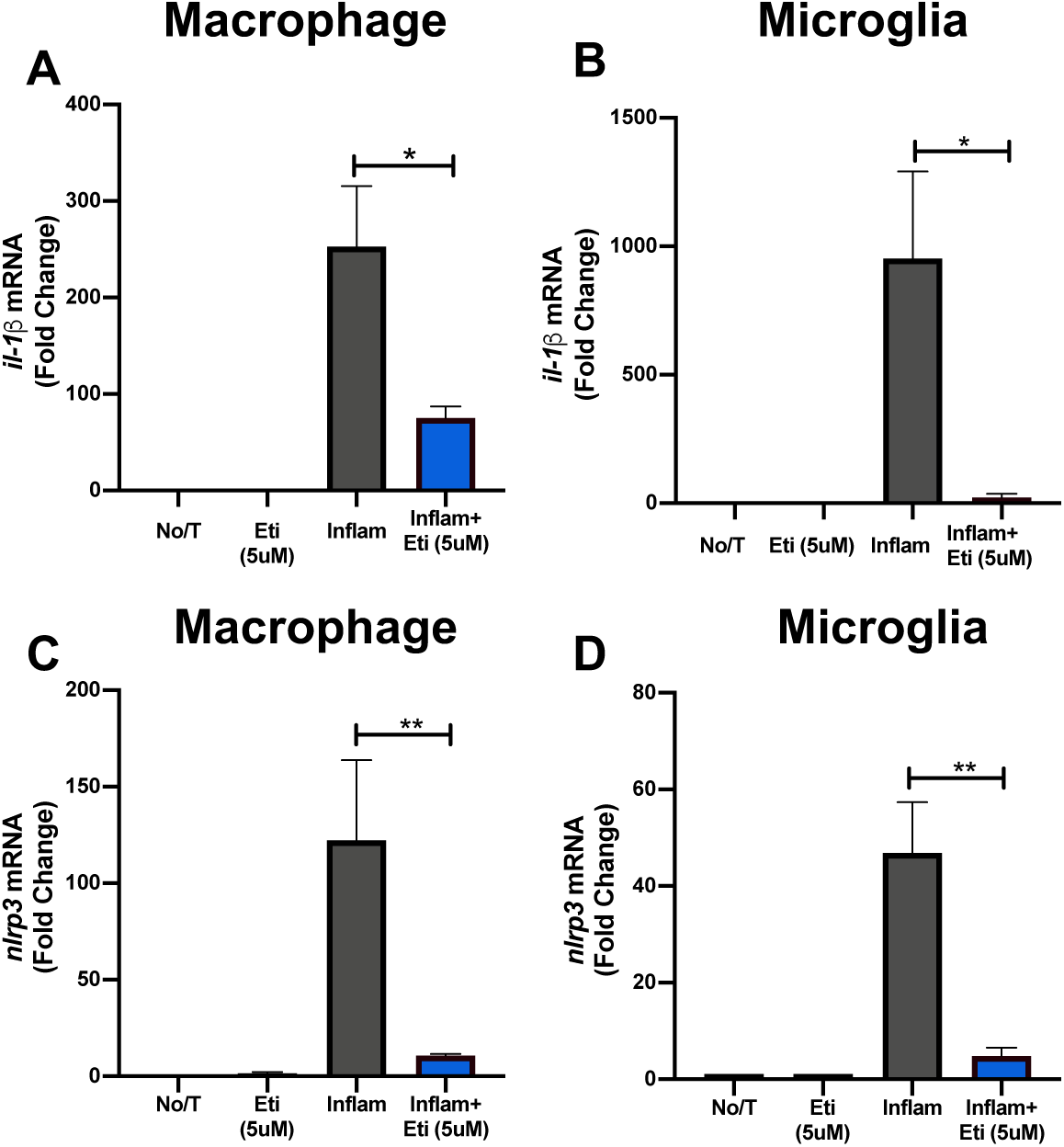
Inflammasome-associated gene expression is decreased in primary mouse macrophages and microglia following pre-treatment with etifoxine (5uM) under inflammasome-activating conditions. RT-qPCR was performed using RNA isolated from primary mouse macrophages and microglia (n=4) that were untreated, treated with etifoxine (5uM) (eti), inflammasome activated (inflam), or pre-treated with etifoxine (5uM) under inflammasome-activating conditions. (A) Fold change of *il-1β* mRNA in macrophages was significantly decreased when pre-treated with etifoxine under inflammasome-activating conditions (73.52±5.42) compared to the inflammasome-only control (252.72±12.81). (B) Fold change of *il-1β* mRNA in microglia was significantly decreased when pre-treated with etifoxine under inflammasome-activating conditions (22.21±5.84) compared to the inflammasome-only control (1147±92.54). (C) Fold change of *nlrp3* mRNA in macrophages was significantly decreased when pre-treated with etifoxine under inflammasome-activating conditions (10.13±1.69) compared to the inflammasome-only control (122.28±26.05). (D) Fold change of *nlrp3* mRNA in microglia was significantly decreased when pre-treated with etifoxine under inflammasome-activating conditions (4.86±1.45) compared to the inflammasome-only control (53.22±13.62). All fold changes were calculated by using the 2^-ΔΔCT^ method. Results are displayed as mean ± SEM. One-way analysis of variance with Tukey’s post hoc test was used to determine group differences. \**p*<0.05, \*\**p*<0.01.

### Pre-treatment with etifoxine decreases inflammasome-associated gene and protein expression in activated human-derived primary macrophages

There was a 128-fold decrease in *il-1β* (Figure 6A), 1.5-fold decrease in *nlrp3* (Figure 6B), and 28-fold decrease in *tnf⍺-ip3* (Figure 6C) expression in murine macrophages pre-treated with etifoxine (50uM) under inflammasome-activating conditions compared to the inflammasome-only controls. Western blotting for NLRP3 protein confirmed a decrease in expression with etifoxine pre-treatment (Figure 6D).

**Figure 6:**
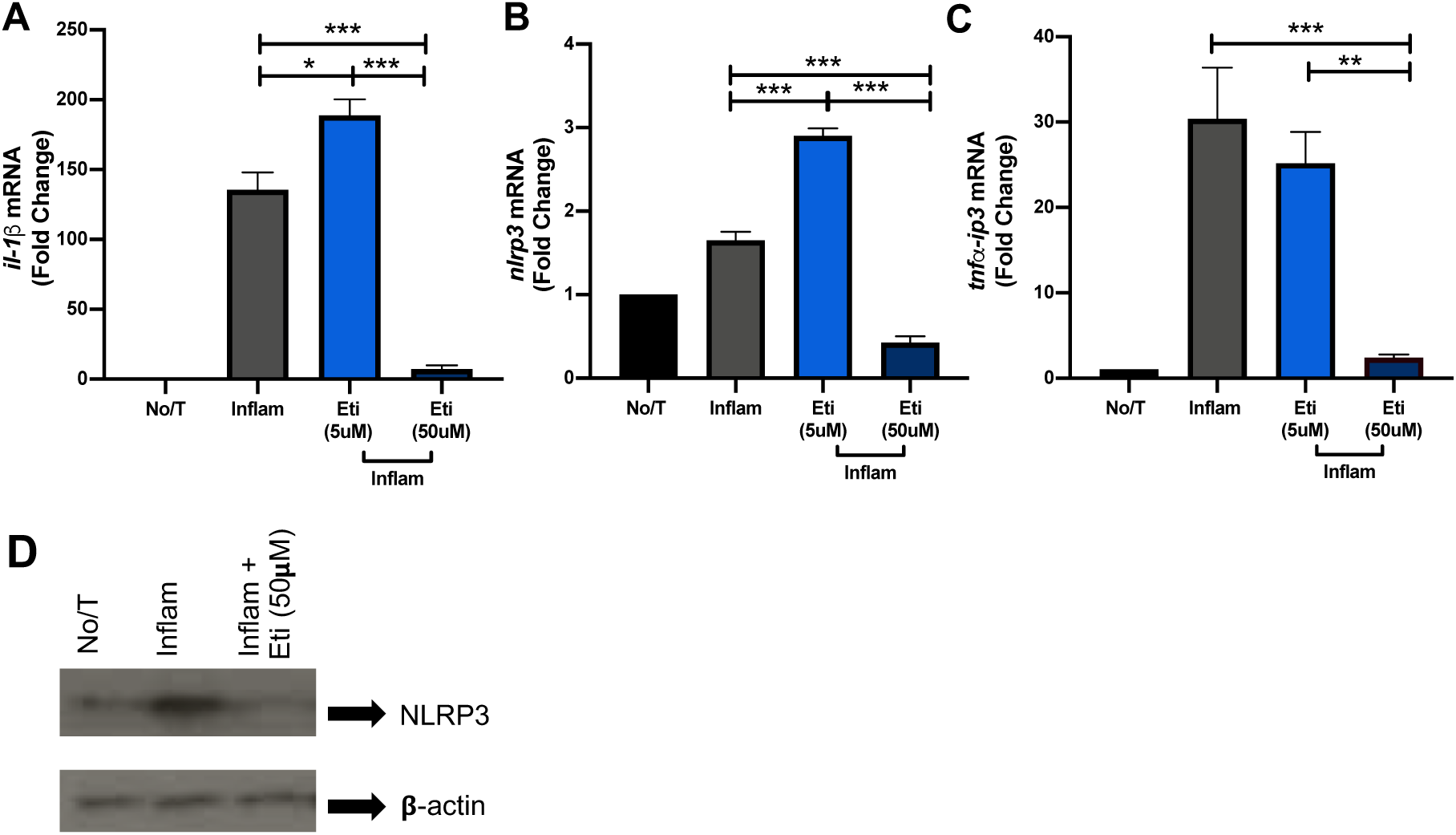
Inflammasome-associated gene expression is decreased when primary human macrophages are pre-treated with 50uM etifoxine under inflammatory conditions. RT-qPCR was performed on primary human macrophages (n=4) that were untreated, inflammasome-activated (inflam), and pre-treated with etifoxine (5uM, 50uM) (eti) under inflammasome-activating conditions. Western blotting for NLRP3 was also performed on primary human macrophages (n=) that were untreated, inflammasome-activated (inflam), and pre-treated with etifoxine (50uM) (eti) under inflammasome-activating conditions. Protein loading was normalized to relative β-actin. Fold change of *il-1β* mRNA when pre-treated with etifoxine (5uM, 50uM) under inflammasome-activating conditions (178.08±13.26, 9.26±2.21) compared to the inflammasome-only control (143.17±15.12). (B) Fold change of *nlrp3* mRNA when pre-treated with etifoxine (5uM, 50uM) under inflammasome-activating conditions (2.9±0.43, 0.43±0.06) compared to the inflammasome-only control (1.65±0.13). (C) Fold change of *tnf⍺-ip3* mRNA when pre-treated with etifoxine (5uM, 50uM) under inflammasome-activating conditions (25.18±2.46, 2.40±0.19) compared to the inflammasome-only control (30.37±3.06). (D) Pre-treatment with etifoxine (50uM) significantly decreased NLRP3 protein expression under inflammasome-activating conditions when compared to the inflammasome-only control. All fold changes were calculated by using the 2^-ΔΔCT^ method. Results are displayed as mean ± SEM. One-way analysis of variance with Tukey’s post hoc test was used to determine group differences. \**p*<0.05, \*\**p*<0.01, \*\*\**p*<0.001.

### Pre-treatment with etifoxine significantly alters inflammasome-associated gene expression in primary mouse macrophages

To further explore additional genes and/or inflammasome-related pathways that may be altered as a result of etifoxine pre-treatment in primary mouse macrophages (other than NLRP3), a Qiagen RT^2^ qPCR array containing 86 inflammasome-specific genes was performed. The conditions compared were inflammasome only vs. no treatment (Figure 7A), etifoxine (5uM) under inflammasome activating conditions vs. no treatment (Figure 7B), and etifoxine (5uM) under inflammasome activating conditions vs. inflammasome only (Figure 7C). Full data is presented in Tables 1-3.

**Figure 7:**
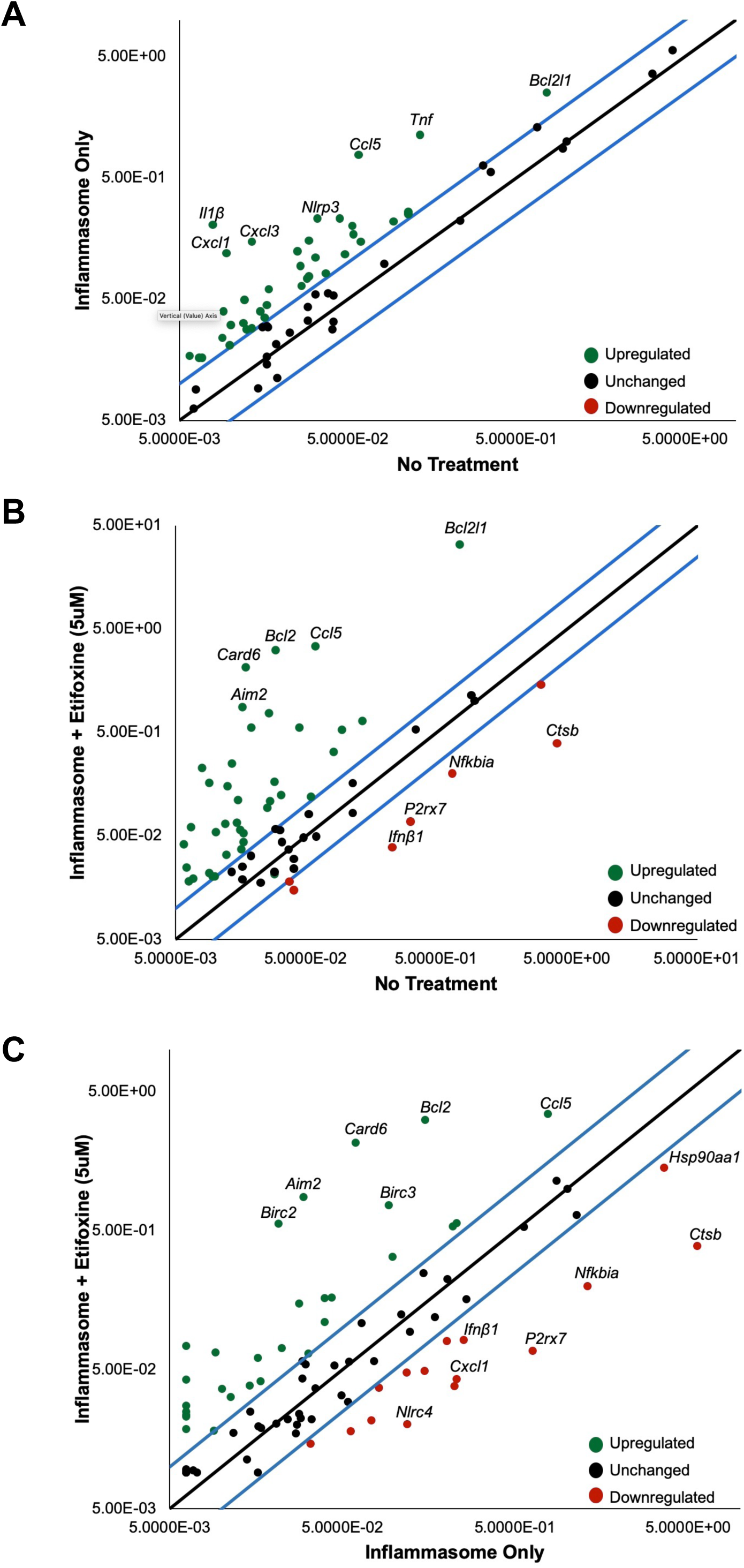
Scatterplots displaying gene expression when comparing inflammasome-activated primary mouse macrophages to untreated control and etifoxine treatment. A Qiagen data analysis web-based program was used to determine the fold regulation by using the 2^-ΔΔCT^ method. Initially the ΔC_T_ is calculated between the genes of interest and the reference genes, which is then followed by the ΔΔC_T_ and fold change calculations between inflammasome-only & no treatment (A), inflammasome + etifoxine & no treatment (B), and inflammasome + etifoxine & inflammasome only (C). (n=3/group). Points are plotted as log_10_ values. The scatterplot compares the normalized gene expression between the test group and the control group, the black line represents unchanged (<2-fold) gene expression and the blue lines represent the >2-fold change threshold. Upregulated genes are colored in green, unchanged genes are colored in black, and downregulated genes are colored in red. Individual points in the upper left and lower right quadrants are up- and downregulated by >2-fold in the inflammasome-only group compared to the control group. The compared groups had a biological n=3. Genes with the greatest increase in fold regulation have been labelled.

### SPMS patient-derived monocytes display an increased susceptibility to LPS treatment compared to healthy controls, while pre-treatment with etifoxine decreases IL-1β secretion in activated SPMS patient-derived PBMCs

To further investigate the potential involvement of inflammasome activation in relation to the progressive nature of MS, we sought to determine the effect that LPS activation had on pro-inflammatory cytokine secretion *in vitro.* Healthy control and SPMS patient-derived monocytes were treated with LPS; LPS treatment alone induces inflammasome activation in monocytes(50). IL-1β (Figure 8A) and TNF (Figure 8C) expression were significantly increased in SPMS patients when compared to healthy controls; IL-6 (Figure 8B) was unchanged. This showed that monocytes derived from SPMS patients were more sensitive to LPS treatment. To determine if the inhibitory effect of etifoxine is potentially clinically relevant, PBMCs derived from SPMS patients were either untreated, inflammasome activated (LPS and nigericin), and pre-treated with etifoxine (50uM) under inflammasome activating conditions. IL-1β secretion was significantly decreased when PBMCs were pre-treated with etifoxine under inflammasome activating conditions when compared to inflammasome-only controls (Figure 8D).

**Figure 8:**
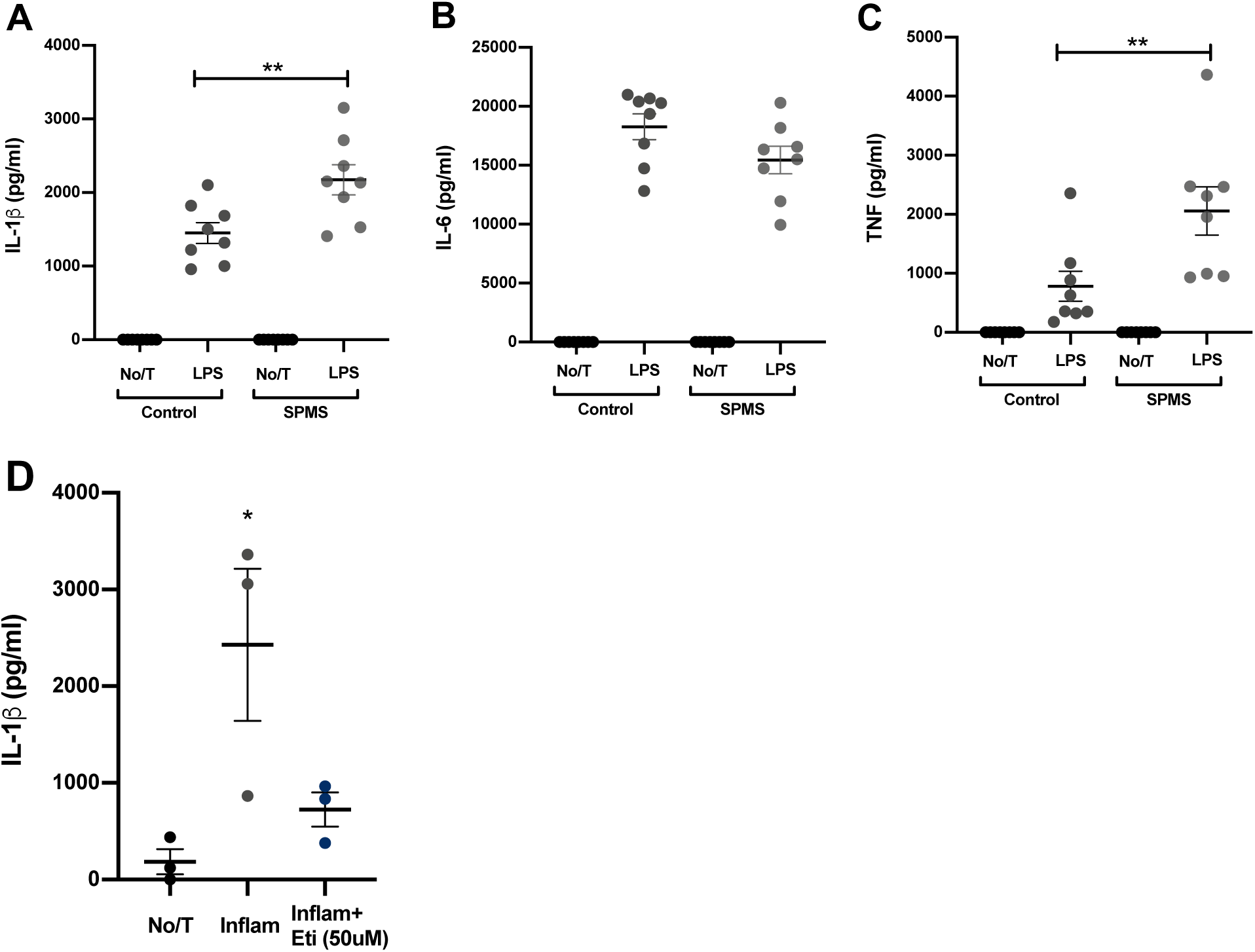
Monocytes derived from SPMS patients display an increased sensitivity to LPS treatment when compared to monocytes derived from age and sex matched healthy controls while pre-treatment with etifoxine (50uM) decreases IL-1β secretion in SPMS patient-derived PBMCs under inflammasome activating conditions. ELISAs were conducted in monocytes derived from healthy controls (n=6/group) and SPMS patients (n=6/group). The control and SPMS monocytes were either untreated or treated with LPS. (A) ELISA comparing IL-1β secretion in monocytes from healthy controls and SPMS patients. IL-1β secretion was significantly increased in SPMS patients (2174pg/mL±204.4) when compared to healthy controls (1451pg/mL±142.3). (B) ELISA comparing IL-6 secretion in monocytes from healthy controls and SPMS patients. There was no significant difference in IL-6 secretion in SPMS patients (15441pg/mL±1164) compared to healthy controls (18266pg/mL±1095). (C) ELISA comparing TNF secretion in monocytes from healthy controls and SPMS patients. TNF secretion was significantly increased in SPMS patients (2055pg/mL±408.8) when compared to healthy controls (781.5pg/mL±254.1). (D) An IL-1β ELISA was conducted on PBMCs derived from SPMS patients (n=3). PBMCs were untreated, inflammasome activated (inflam), and pre-treated with etifoxine (50uM) (eti) under inflammasome activating conditions. IL-1 secretion was significantly decreased when pre-treated with etifoxine (50uM) under inflammasome activating conditions (725.6pg/mL±77.5) when compared to inflammasome-only control (2428pg/mL±786.7). Results are displayed as mean ± SEM. One-way analysis of variance with Tukey’s post hoc test was used to determine group differences. \**p*<0.05. \*\**p*<0.01

## DISCUSSION

In this study, we investigated the ability of etifoxine, a small-molecule TSPO ligand, to influence inflammasome activation in murine- and human-derived myeloid cells in the pathological context of MS. To further elucidate the inflammasome-associated mechanism of action for etifoxine, we investigated whether IL-1β secretion in myeloid cells was altered when cells were pre-treated with etifoxine under inflammasome-activating conditions. Etifoxine significantly decreased IL-1β release in both murine macrophages and microglia, which supports prior evidence that etifoxine possesses an anti-inflammatory effect in murine myeloid cells (25, 51, 52). Similar results were also observed using human myeloid cells. We then conducted a time course-dependent assay where etifoxine was used as: 1) a pre-treatment, 2) in combination with LPS, or 3) in combination with nigericin. Results of these experiments confirmed that etifoxine was likely exerting its anti-inflammatory effect when used as a pre-treatment and in combination with LPS, since etifoxine had no effect on IL-1β secretion when used at the stage of nigericin treatment. These findings suggest that etifoxine is likely working via a mechanism that is upstream of inflammasome assembly and/or activation, perhaps by inhibiting components of the LPS-driven TLR4 pathway (53, 54).

In addition to altering IL-1β secretion, pre-treatment with etifoxine also inhibited the expression of inflammasome-related genes *in vitro* in both mouse and human myeloid cells. Pre-treatment with etifoxine decreased *nlrp3* and *il-1β* mRNA expression in both species under inflammasome-activating conditions, further validating the hypothesis that etifoxine is inhibiting inflammasome activation. Furthermore, to confirm that decreased levels of cytokines were not merely due to a potential cytotoxic effect of etifoxine, cytotoxicity and cell viability assays were performed in both murine- and human-derived macrophages. A wide dose range of etifoxine was used and no effect on cytotoxicity or cell viability were observed.

As stated previously, etifoxine has two previously identified binding partners: TSPO and GABA_A_ receptors. GABAergic neurotransmission is highly associated with neurosteroid synthesis; however, microglia are unable to produce steroids due to an absence of the CYP11A1 enzyme (25, 32, 55, 56). Herein, we provide evidence that excludes GABA_A_ receptors as a possible mechanism of action for etifoxine. In the presence of GABA_A_ inhibition, etifoxine was still potently able to significantly decrease inflammasome activity and IL-1β production, further suggesting the presence of a novel mechanism of action for etifoxine in terms of inhibiting inflammation.

*In vitro*, we have demonstrated that etifoxine decreases pro-inflammatory cytokine secretion and inhibits inflammasome activation in myeloid cells derived from healthy individuals. However, to further implicate etifoxine as a potential therapetic in the pathological context, we further investigated etifoxine in relation to MS disease pathology. There is increasing evidence that supports a role for NLRP3 and inflammasome-related genes in SPMS disease progression (11, 13, 20, 39, 51), especially when considering the increase in microglia activation that characterizes SPMS. To determine the potential for etifoxine to influence inflammasome activity in SPMS-derived cells, we utilized our resources to conduct multiple experiments in SPMS-patient derived monocytes (20, 21, 57–59). In this series of experiments, LPS treatment was used to activate SPMS-patient derived monocytes *in vitro* (60), which allowed us to observe a variablity in cytokine secretion between healthy controls and SPMS patients. In cells derived from SPMS patients, LPS treatment resulted in a significant increase in IL-1β and TNF secretion compared to age and sex-matched healthy controls.

Lastly, we wanted to determine if pre-treatment with etifoxine is capable of inducing an anti-inflammatory effect in SPMS patient-derived PBMCs under inflammasome activating conditions. In MS, inflammasomes have been shown to be associated with susceptibilty, disease severity, and disease progression; it has also been suggested that inhibiting inflammasome activation may act as a form of therapy by reducing neuroinflammation (51, 52, 61–63). Our results demonstrate that pre-treatment with etifoxine has an inhibitory effect on IL-1β expression in SPMS-patient derived PBMCs under inflammasome activating conditions. This finding can be utilized to further explore the therapeutic potential of etifoxine in SPMS.

## DECLARATIONS

### Ethics approval and consent to participate

All studies that involved human samples followed Canadian Institute of Health Research (CIHR) guidelines and institutional review board approval at Memorial University of Newfoundland (Health Research Ethics Authority). All studies that involved animals were conducted following institutional ethics approval from Memorial University’s animal care committee following recommended guideline from the Canadian Council for Animal Care

### Consent for Publication

Not applicable

### Availability of data and materials

The datasets used and/or analyzed during the current study are available from the corresponding author on reasonable request

### Authors’ contributions

JMO designed and performed experiments, analyzed data, and wrote the initial manuscript. JBW assisted in developing and optimizing in vitro assays. DRO & PMM assisted in designing experiments, interpreting data, and reviewing the manuscript. CSM secured funding for this project, conceived initial project idea, designed and performed experiments, analyzed data, interpreted results, and prepared and reviewed the finalized manuscript.

### Author’s information

Not applicable

## Acknowledgements

The authors wish to acknowledge and thank all study participants involved in this research.

