## Supplemental Figures and Tables for "Etifoxine inhibits NLRP3 inflammasome activity in human and murine myeloid cells"

#### Supplemental Figure 1.

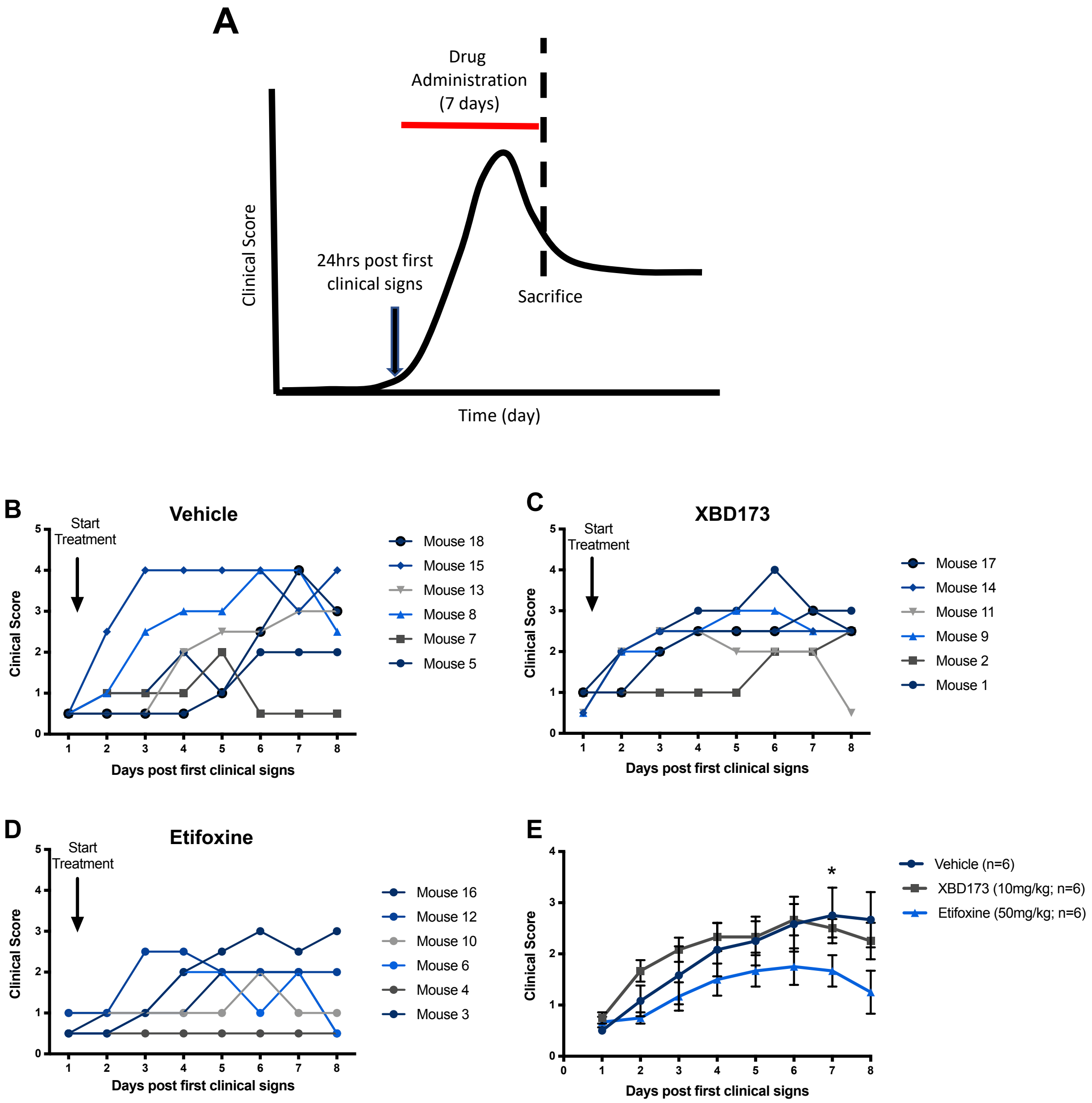


**Supplemental Figure 1**: (A) In EAE mice, vehicle, XBD173, or etifoxine treatment was initiated 24 hours after the first clinical signs were observed and continued for 7 days. (B) Clinical score for the vehicle treatment group (n=6) peaked at a mean score of 2.75 on day 7. (C) Clinical score for the XBD173 treatment group (n=6) peaked at mean score of 2.66 on day 6. (D) Clinical score for the etifoxine treatment group (n=6) peaked at a mean score of 1.75 on day 6 and dropped to a mean score of 1.25 on day 8. (E) Clinical score of each treatment group plotted together. Repeated Measures one-way analysis of variance was used to determine group differences between vehicle and XBD173 (*p*=0.53), vehicle and etifoxine (*p*<0.05), and etifoxine and XBD173 (*p*<0.01).

**Supplemental Figure 2.**

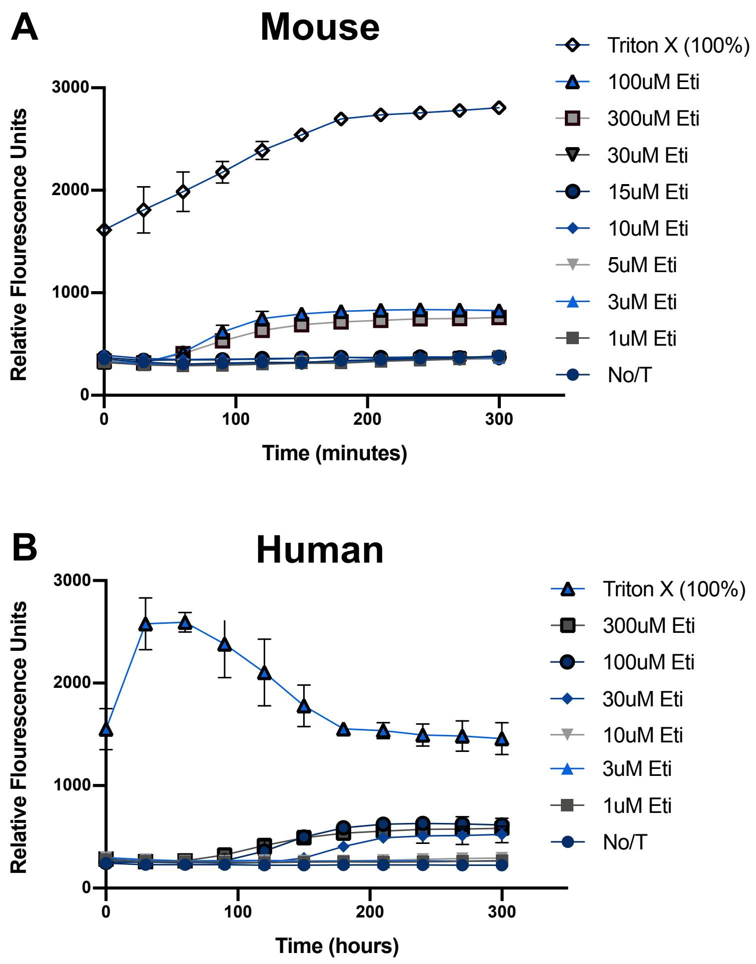


**Supplemental Figure 2: In human and mouse macrophages, etifoxine is not cytotoxic at doses that block IL-1β secretion.** Cytotoxicity assays were conducted in mouse and human primary macrophages (n=2) and all treatments occurred in the absence of inflammasome-activating conditions. The assays were conducted in real-time using the Cytation$^{TM}$ 5 Cell Imaging Multi-Mode Reader for 5 hours. (A) Primary mouse macrophages were untreated, treated with etifoxine (1uM-300uM) (eti), and treated with Triton X. The Triton X condition peaked and plateaued at approximately 2700 MFU, the etifoxine (100uM, 300uM) conditions peaked at 800 MFU, and the lower doses (1uM-30uM) were consistent with the untreated control. (B) Human macrophages were untreated, treated with etifoxine (1uM-300uM), and treated with Triton X. The Triton X condition peaked at approximately 2600 MFU and steadily decreased to 1500 MFU, the etifoxine (30uM-300uM) conditions peaked at 650 MFU, and the lower doses (1uM-10uM) were consistent with the untreated control.

#### Supplemental Figure 3.

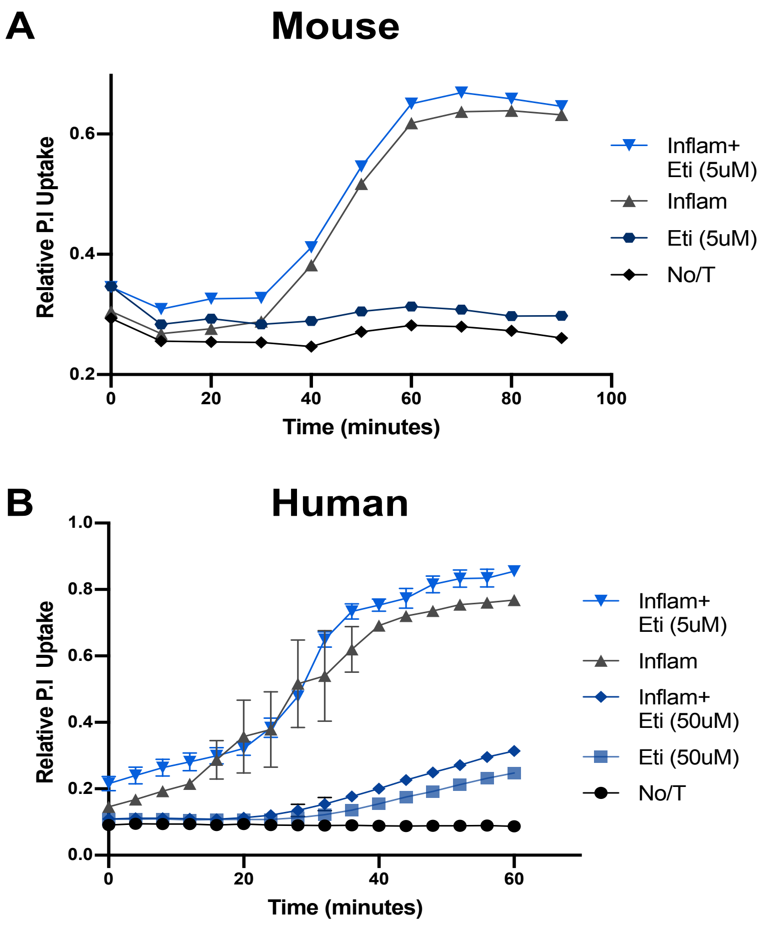


**Supplemental Figure 3: Propidium iodide uptake is unchanged in primary mouse macrophages when pre-treated with etifoxine (5uM), and decreased in primary human macrophages when pre-treated with etifoxine (50uM) under inflammasome-activating conditions.** Propidium iodide uptake assays were conducted in primary mouse macrophages (n=1) and primary human macrophages (n=2). The assays were conducted in real-time using the Cytation$^{TM}$ 5 Cell Imaging Multi-Mode Reader for 1-1.5 hours. (A) Primary mouse macrophages were untreated, treated with etifoxine (5uM) (eti), inflammasome activated (inflam), and pre-treated with etifoxine (5uM) under inflammasome-activating conditions. PI uptake in macrophages that were pre-treated with etifoxine under inflammasome-activating conditions did not differ from the inflammasome-only control, there was also no difference between the etifoxine condition and the untreated control. (B) Primary human macrophages were untreated, treated with etifoxine (50uM), inflammasome activated, and pre-treated etifoxine (5uM,50uM) under inflammasome activating conditions. PI uptake was reduced when macrophages were pre-treated with etifoxine (50uM) under inflammasome-activating conditions compared to the inflammasome-only control and did not differ compared to the untreated control.

### Supplemental Figure 4.

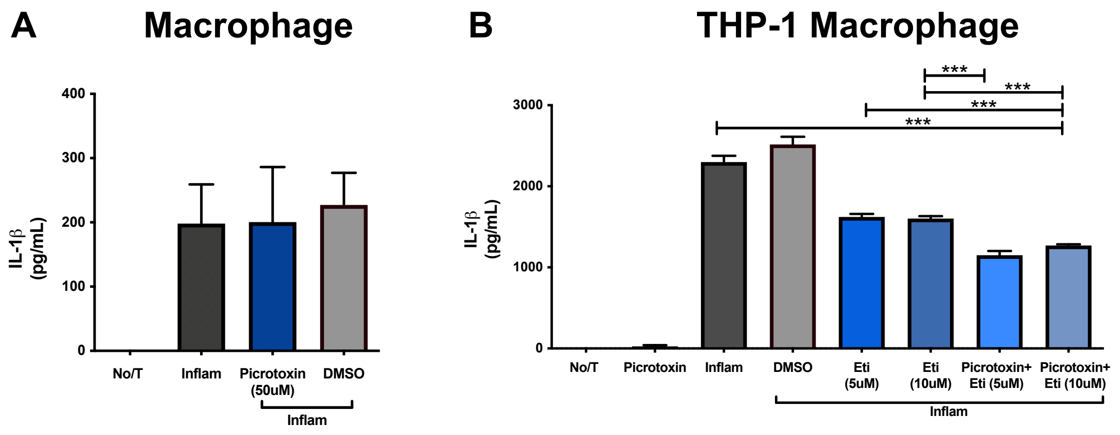


**Supplemental Figure 4: IL-1β secretion is unchanged when primary mouse BMDMs and human THP-1 macrophages are pre-treated with picrotoxin.** IL-1β ELISAs were conducted in primary mouse macrophages (n=3) and human THP-1 macrophages (n=6). (A) Primary mouse macrophages were untreated, inflammasome activated (inflam), and pre-treated with either picrotoxin (50uM) or DMSO under inflammasome-activating conditions. IL-1β secretion did not differ when comparing the pre-treatment with picrotoxin under inflammasome-activating conditions (200pg/mL±85.91) to the inflammasome-only control (198pg/mL±60.89). (B) THP-1 macrophages were untreated, treated with picrotoxin (50uM), inflammasome activated, and pre-treated with either DMSO, etifoxine alone (5uM, 10uM) (eti), or picrotoxin (50uM) and etifoxine (5uM, 10uM) under inflammasome-activating conditions. Picrotoxin alone did not affect IL-1β secretion when compared to the untreated control. IL-1β secretion was decreased when pre-treated with etifoxine (5uM, 10uM) alone (1621pg/mL±38.15, 1602pg/mL±130.28) and with the addition of picrotoxin (1149pg/mL±52.27, 1269pg/mL±15.97) under inflammasome-activating conditions compared to the inflammasome-only control. Results are displayed as mean ± SEM. One-way analysis of variance with Tukey’s post hoc test was used to determine group differences. ****p*<0.001.

**Supplemental Figure 5.**

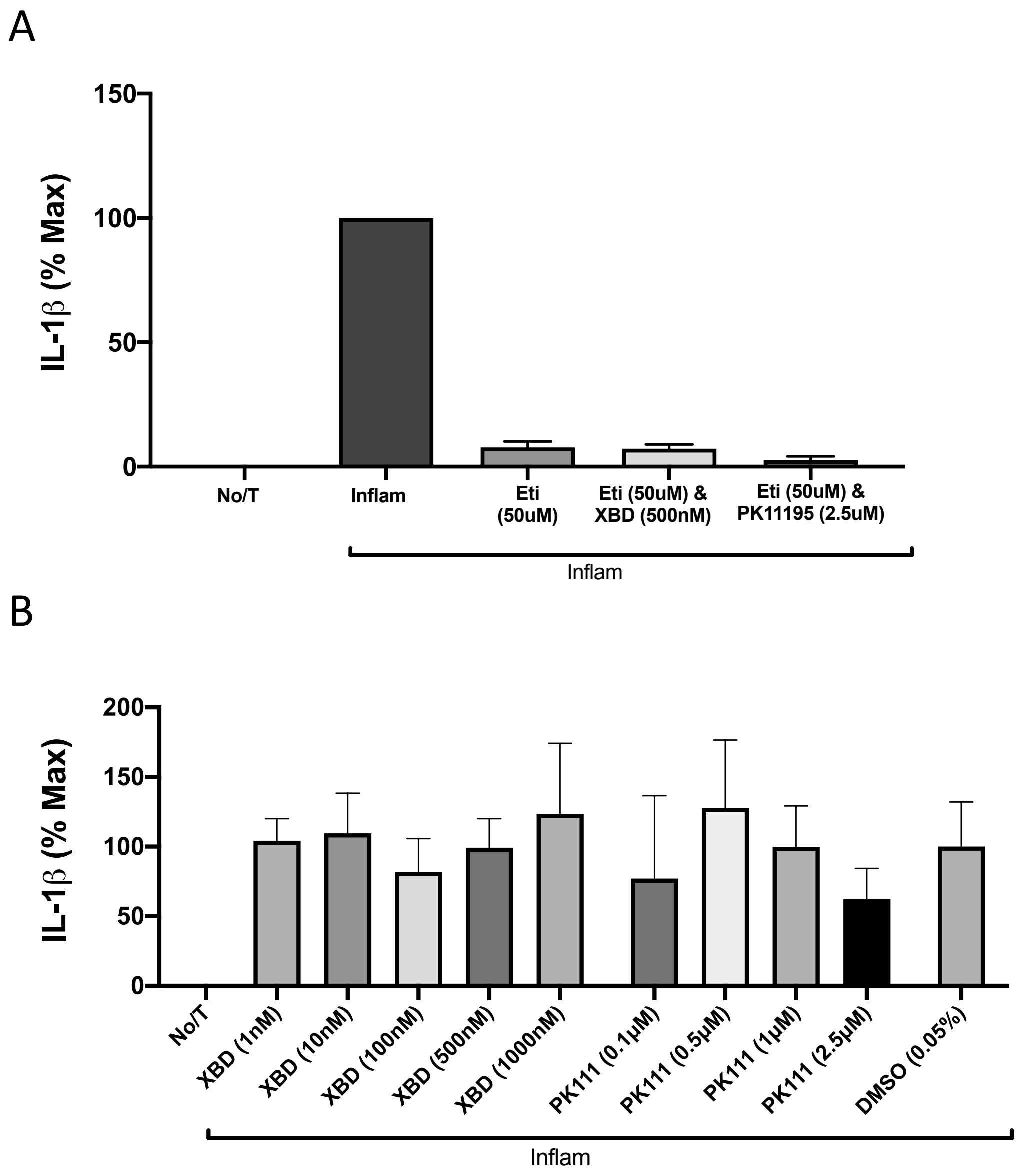


**Supplemental Figure 5: IL-1β secretion is unchanged when primary macrophages are simultaneously treated with etifoxine and established TSPO ligands or with TSPO ligands alone.** IL-1β ELISAs were conducted on supernatants from primary human macrophages (n=2). Primary macrophages were untreated, inflammasome activated (inflam), or pre-treated with either etifoxine (50uM) alone, etifoxine (50uM) & XBD173 (500nM), or etifoxine (50uM) & PK11195 (2.5uM) for 1hr prior to inflammasome activation. IL-1β secretion did not differ when comparing the etifoxine-treated alone vs. etifoxine in combination with known TSPO ligands. B) IL-1β ELISAs were conducted on supernatants from primary mouse macrophages (n=3). Primary macrophages were untreated, inflammasome activated (inflam), or pre-treated with either XBD173 (1-1000nM) or PK11195 (0.1-2.5uM) for 1hr prior to inflammasome activation. IL-1β secretion did not differ between treatment groups. Results are displayed as mean ± SEM.

**Table 1: Gene Regulation in inflammasome activated primary mouse macrophages when compared to untreated control. Upregulated genes (>2-fold change) are colored in green and unchanged genes (<2-fold change) are colored in black**

| Gene | Regulation | Gene | Regulation |
|---|---|---|---|
| *Il1β* | 25.34 | *Fadd* | 2.47 |
| *Ccl5* | 13.21 | *Mapk13* | 2.46 |
| *Cxcl1* | 12.37 | *Casp1* | 2.44 |
| *Cxcl3* | 10.91 | *Tab1* | 2.39 |
| *Tnf* | 8.44 | *Tnfsf14* | 2.38 |
| *Il6* | 8.31 | *Tab2* | 2.31 |
| *Nlrp3* | 6.9 | *Rela* | 2.26 |
| *Il12b* | 6.06 | *Ccl7* | 2.23 |
| *Irf4* | 6 | *Mapk3* | 2.18 |
| *Ifnβ1* | 5.19 | *Nfkb1* | 2.18 |
| *Ptgs2* | 5.1 | *Irak1* | 2.16 |
| *Bcl2* | 5.09 | *Nfkbib* | 2.1 |
| *Nlrx1* | 4.84 | *Pycard* | 2.08 |
| *Ccl12* | 4.31 | *P2rx7* | 1.98 |
| *Tnfsf4* | 4.31 | *Nfkbia* | 1.95 |
| *Ciita* | 4.03 | *Nlrc5* | 1.87 |
| *Irf1* | 3.78 | *Aim2* | 1.82 |
| *Myd88* | 3.68 | *Mapk12* | 1.76 |
| *Nlrp1a* | 3.61 | *Nod2* | 1.75 |
| *Ifng* | 3.53 | *Mok* | 1.74 |
| *Il12a* | 3.53 | *Irf3* | 1.71 |
| *Naip1* | 3.53 | *Mapk11* | 1.63 |
| *Nlrp5* | 3.53 | *Hsp90b1* | 1.59 |
| *Nlrp6* | 3.53 | *Txnip* | 1.51 |
| *Nlrp9b* | 3.53 | *Xiap* | 1.47 |
| *Birc3* | 3.52 | *Nlrc4* | 1.43 |
| *Card6* | 3.47 | *Sugt1* | 1.34 |
| *Nlrp4e* | 3.4 | *Ctsb* | 1.33 |
| *Cd40lg* | 3.37 | *Ikbkb* | 1.2 |
| *Ripk2* | 3.37 | *Mapk8* | 1.18 |
| *Bcl2l1* | 3.27 | *Map3k7* | 1.15 |
| *Cflar* | 3.14 | *Birc2* | 1.13 |
| *Pstpip1* | 2.99 | *Hsp90aa1* | 1.12 |
| *Nlrp4b* | 2.95 | *Il18* | 1.03 |
| *Nlrp12* | 2.92 | *Il33* | 1.02 |
| *Naip5* | 2.91 | *Mapk1* | 0.95 |
| *Traf6* | 2.72 | *Pea15a* | 0.92 |
| *Mefv* | 2.64 | *Tirap* | 0.88 |
| *Nod1* | 2.62 | *Panx1* | 0.8 |
| *Casp8* | 2.59 | *Mapk9* | 0.7 |
| *Tnfsf11* | 2.57 | *Casp12* | 0.63 |
| *Chuk* | 2.55 | *Ikbkg* | 0.59 |

**Table 2: Gene Regulation in primary mouse macrophages pre-treated with etifoxine (5µM) under inflammasome-activating conditions when compared to untreated control. Upregulated genes (>2-fold change) are colored in green, downregulated genes are colored in red, and unchanged genes (<2-fold change) are colored in black**

| Gene | Regulation | Gene | Regulation |
|---|---|---|---|
| *Card6* | 124.78 | *Irf4* | 3.37 |
| *Bcl2* | 106.16 | *Traf6* | 3.25 |
| *Ccl5* | 59.14 | *Myd88* | 2.98 |
| *Aim2* | 53.28 | *Nlrc4* | 2.91 |
| *Bcl2l1* | 43.19 | *Tnfsf14* | 2.85 |
| *Naip1* | 41.97 | *Ciita* | 2.67 |
| *Birc2* | 29.88 | *Nod2* | 2.59 |
| *Birc3* | 29.11 | *Mapk12* | 2.58 |
| *Il1β* | 28.16 | *Mok* | 2.56 |
| *Nlrp5* | 24.22 | *Mefv* | 2.42 |
| *Il12b* | 19.35 | *Irak1* | 2.29 |
| *Cxcl3* | 18.39 | *Cflar* | 2.2 |
| *Ccl12* | 17.82 | *Cxcl1* | 2.11 |
| *Nlrp6* | 15.65 | *Mapk11* | 2.07 |
| *Ifng* | 13.8 | *Pycard* | 2.06 |
| *Il12a* | 13.11 | *Tnfsf11* | 1.94 |
| *Nlrp1a* | 13.07 | *Irf3* | 1.79 |
| *Cd40lg* | 12.85 | *Ikbkg* | 1.67 |
| *Ptgs2* | 12.69 | *Nfkbib* | 1.65 |
| *Ccl7* | 11.95 | *Hsp90b1* | 1.51 |
| *Mapk13* | 9.29 | *Irf1* | 1.51 |
| *Nlrp12* | 8.78 | *Tirap* | 1.5 |
| *Casp8* | 7.25 | *Rela* | 1.41 |
| *Naip5* | 7.18 | *Nlrp3* | 1.3 |
| *Tnfsf4* | 6.48 | *Pea15a* | 1.21 |
| *Il6* | 6.12 | *Il18* | 1.16 |
| *Txnip* | 5.77 | *Casp1* | 0.99 |
| *Tab2* | 5.74 | *Mapk3* | 0.99 |
| *Nlrp9b* | 5.47 | *Fadd* | 0.82 |
| *Nod1* | 5.4 | *Mapk8* | 0.77 |
| *Pstpip1* | 5.37 | *Map3k7* | 0.76 |
| *Ifnβ1* | 5.34 | *Chuk* | 0.75 |
| *Tnf* | 4.84 | *Sugt1* | 0.72 |
| *Nlrp4e* | 4.71 | *Nfkb1* | 0.71 |
| *Casp12* | 4.52 | *Mapk9* | 0.6 |
| *Nlrp4b* | 4.32 | *Ctsb* | 10.88 |
| *Il33* | 4.11 | *Mapk1* | 6.11 |
| *Tab1* | 4.03 | *P2rx7* | 4.65 |
| *Ikbkb* | 3.96 | *Nfkbia* | 3.34 |
| *Ripk2* | 3.85 | *Panx1* | 2.77 |
| *Nlrx1* | 3.72 | *Hsp90aa1* | 2.27 |
| *Nlrc5* | 3.68 | *Xiap* | 2.11 |

**Table 3: Gene Regulation in primary mouse macrophages pre-treated with etifoxine (5μM) under inflammasome-activating conditions when compared to inflammasome-only control. Upregulated genes (>2-fold change) are colored in green, downregulated genes are colored in red, and unchanged genes (<2-fold change) are colored in black**

| Gene | Regulation | Gene | Regulation |
|---|---|---|---|
| *Card6* | 36 | *Nlrp4e* | 1.38 |
| *Aim2* | 29.33 | *Pea15a* | 1.32 |
| *Birc2* | 26.46 | *Mapk11* | 1.27 |
| *Bcl2* | 20.85 | *Tnfsf14* | 1.2 |
| *Bcl2l1* | 13.21 | *Traf6* | 1.2 |
| *Naip1* | 11.88 | *Ripk2* | 1.14 |
| *Birc3* | 8.27 | *Il18* | 1.12 |
| *Casp12* | 7.23 | *Il1β* | 1.11 |
| *Nlrp5* | 6.86 | *Irak1* | 1.06 |
| *Ccl7* | 5.37 | *Irf3* | 1.05 |
| *Ccl5* | 4.48 | *Pycard* | 0.99 |
| *Nlrp6* | 4.43 | *Hsp90b1* | 0.95 |
| *Ccl12* | 4.13 | *Mefv* | 0.92 |
| *Il33* | 4.01 | *Mapk9* | 0.86 |
| *Cd40lg* | 3.82 | *Nfkbib* | 0.78 |
| *Txnip* | 3.82 | *Nlrx1* | 0.77 |
| *Mapk13* | 3.77 | *Tnfsf11* | 0.76 |
| *Il12a* | 3.71 | *Il6* | 0.74 |
| *Nlrp1a* | 3.62 | *Cflar* | 0.7 |
| *Ikbkb* | 3.31 | *Map3k7* | 0.67 |
| *Il12b* | 3.19 | *Ciita* | 0.66 |
| *Nlrp12* | 3.01 | *Mapk8* | 0.65 |
| *Ikbkg* | 2.84 | *Myd88* | 0.65 |
| *Casp8* | 2.79 | *Rela* | 0.62 |
| *Ifng* | 2.66 | *Tnf* | 0.57 |
| *Ptgs2* | 2.49 | *Irf4* | 0.56 |
| *Tab2* | 2.48 | *Tab1* | 0.55 |
| *Naip5* | 2.47 | *Ctsb* | 14.5 |
| *Nod1* | 2.06 | *P2rx7* | 9.22 |
| *Nlrc4* | 2.03 | *Nfkbia* | 6.5 |
| *Nlrc5* | 1.97 | *Cxcl1* | 5.85 |
| *Pstpip1* | 1.8 | *Mapk1* | 5.82 |
| *Tirap* | 1.7 | *Nlrp3* | 5.3 |
| *Cxcl3* | 1.69 | *Chuk* | 3.39 |
| *Nlrp9b* | 1.55 | *Xiap* | 3.1 |
| *Ifnβ1* | 1.51 | *Nfkb1* | 3.07 |
| *Tnfsf4* | 1.5 | *Fadd* | 3.02 |
| *Nod2* | 1.48 | *Hsp90aa1* | 2.54 |
| *Mapk12* | 1.47 | *Irf1* | 2.5 |
| *Mok* | 1.47 | *Casp1* | 2.46 |
| *Nlrp4b* | 1.47 | *Panx1* | 2.23 |
| *Sugt1* | 1.47 | *Mapk3* | 2.19 |

**Table 4: Patient demographics for the healthy control and SPMS patient-derived monocytes**

| | Control | SPMS |
|---|---|---|
| **Age** | 56±7.2 (41-63) | 53.8±10.0 (43-73) |
| **Sex** | 5♀; 3♂ | 5♀; 3♂ |
| **EDSS** | N/A | 6.5-7.0 |
| ***no DMT use** | | |
